# S1PR1 Regulates Lymphatic Valve Development And Prevents Ileitis-Independent Tertiary Lymphoid Organ Formation

**DOI:** 10.1101/2024.09.30.615915

**Authors:** Xin Geng, Lijuan Chen, Zoheb Ahmed, Guilherme Pedron Formigari, Yenchun Ho, Ilaria Del Gaudio, Marcella Neves Datilo, Zheila J Azartash-Namin, Xindi Shan, Ravi Shankar Keshari, Hong Chen, Florea Lupu, Lijun Xia, Gwendalyn J. Randolph, Scott D Zawieja, Eric Camerer, Michael J Davis, R. Sathish Srinivasan

## Abstract

Lymphatic vessels function throughout the body to drain interstitial fluids. Efficient lymph flow is ensured by lymphatic valves (LVs). However, the mechanisms that regulate LV development are incompletely understood. Here, we show that the deletion of the GPCR sphingosine 1-phosphate receptor-1 (S1PR1) from lymphatic endothelial cells (LECs) results in fewer LVs. Interestingly, LVs that remained in the terminal-ileum draining lymphatic vessels were specifically dysfunctional, and tertiary lymphoid organs (TLOs) formed in this location. TLOs in the terminal ileum are associated with ileitis in humans and mice. However, mice lacking S1PR1 did not develop obvious characteristics of ileitis. Sphingosine kinases 1 and 2 (SPHK1/2) are required for the synthesis of S1P, the ligand of S1PR1. Mice that lack *Sphk1/2* in LECs recapitulate the LV and TLO phenotypes of mice that lack S1PR1. Mechanistically, S1PR1 regulates shear stress signaling and the expression of the valve-regulatory molecules FOXC2 and connexin-37. Importantly, *Foxc2^+/-^* mice, a model for lymphedema-distichiasis syndrome, also develop TLOs in the terminal ileum. Thus, we have discovered S1PR1 as a previously unknown regulator of LV and TLO development. We also suggest that TLOs are a sign of subclinical inflammation that can form due to lymphatic disorders in the absence of ileitis.

## INTRODUCTION

Tertiary lymphoid structures, also known as tertiary lymphoid organs (TLOs) are lymph node-like structures that form under pathological conditions such as in certain tumors, rejected transplant tissues, lungs of COPD or influenza-infected patients, adipose tissue of obese individuals and in tissues afflicted with autoimmune diseases such as Hashimoto thyroiditis, type 1 diabetes, multiple sclerosis, rheumatoid arthritis and inflammatory bowel disease (IBD) (1). TLOs contain B cells, T cells, macrophages, stromal cells such as fibroblastic reticular cells (FRCs) and follicular dendritic cells, and high endothelial venules (HEVs). Infectious agents such as *Helicobacter pylori*, *Helicobacter hepaticus, Borrelia burgdorferi* and *Salmonella enterica* can also promote TLO development (2–4). TLOs are thought to function as sites of localized immune response against microbes and parasites, tumor antigens, grafted tissues, and auto antigens. The presence of TLOs is associated with better prognosis in the context of tumor progression and immune checkpoint therapy (2, 5–8). In contrast, TLOs are thought to aggravate tissue damage in autoimmune diseases and contribute to transplanted tissue rejection (3, 9–12). TLOs can progress into lymphomas as in the context of *Helicobacter pylori* infections (13, 14). TLOs that develop in adipose tissues during obesity can reduce lipolysis and insulin sensitivity (15, 16). Thus, modulation of TLO formation may be beneficial under various pathological conditions. To achieve such modulation, it is critical to obtain a better understanding of how they form.

TLOs share structural similarities with secondary lymphoid organs such as lymph nodes (1). Lymph node development is initiated by the extravasation of hematopoietic-derived lymphoid tissue initiator (LTi) cells at specialized vein junctions that are devoid of smooth muscle cell coverage (17). Nearby lymphatic vessels undergo a cup-shaped morphogenesis to collect LTi and nearby stromal cells, known as lymphoid tissue organizer (LTo) cells. Crosstalk between LTi, LTo, and lymphatic endothelial cells (LECs) promotes the organization and expansion of lymph node primordia. Lymph nodes do not develop properly in mice with defective lymphatic vessels (17–20). Naïve T and B cells enter the mature lymph nodes through HEVs and reside in specific compartments. Afferent lymphatic vessels secrete chemokines, such as CCL21, to recruit antigen loaded DCs from peripheral tissues, which are then transported to lymph nodes where they encounter and activate naïve T and B cells. Efferent lymphatic vessels secrete sphingosine 1-phosphate (S1P) to promote the exit of lymphocytes from the lymph nodes into the lymphatic vessels and then into blood (21–23). Thus, lymphatic vessels are an integral part of lymph node development and function.

TLOs appear to be heterogeneous with respect to the presence or absence of lymphatic vessels. TLOs in adipose tissue were reported to lack lymphatic vessels (24) and TLOs in the lungs developed in the absence of lymphatic vessels or lymphatic drainage (25). However, more recently, TLOs that were connected to dysfunctional mesenteric lymphatic vessels were identified in high fat diet-fed mice and in *Tnf^+/ΔARE^* mice (a model for ileitis) (11, 16, 26). These publications demonstrated that chronic inflammation can result in TLO formation. Despite these findings the relationship between the lymphatic vasculature and TLOs is not fully understood. Whether lymphatic dysfunction can result in TLO formation in the absence of chronic inflammation is also not known.

The lymphatic vasculature consists of lymphatic capillaries, collecting lymphatic vessels, lymphatic valves (LVs) and lymphovenous valves (LVVs) (27). LVVs, LVs and venous valves are structurally similar to each other and express a similar set of regulatory genes (27–30). Vascular valves develop in a stepwise process that involves sensing of oscillatory shear stress (OSS), delamination of cells, cell elongation and collective cell migration (27, 29, 31–36). The GPCR sphingosine 1-phosphate receptor-1 (S1PR1) is an important therapeutic target. Inhibitors of S1PR1 are used to treat inflammatory diseases such as multiple sclerosis and IBD, which are diseases that are likely modulated by the lymphatic vasculature (37, 38). We recently identified S1PR1 as a regulator of LEC junctions, cytoskeleton and lymphangiogenesis (39). In this study we tested whether S1PR1 regulated LV and LVV development and maintenance and serendipitously discovered a previously unknown relationship between S1PR1 signaling, LV development and TLO formation.

## RESULTS

### S1PR1 regulates the development of embryonic lymphatic, lymphovenous and venous valves

Two pairs of LVVs are bilaterally located at the junction of jugular and subclavian veins to regulate lymph return to the blood circulation (28, 29). LVVs are the first vascular valves to develop in mammals. We used the S1PR1 activity reporter mice (“tango” mice, henceforth referred to as S1PR1-GS mice) to test if S1PR1 signaling is active in LVVs (40). In these mice the interaction of S1PR1 with its ligand S1P and the subsequent β-arrestin coupling to the receptor results in the nuclear expression of GFP. By analyzing embryonic day (E) 16.5 S1PR1-GS embryos we determined that S1PR1 signaling was indeed active in LVVs (**Figure 1A**, arrows). S1PR1 activity was also observed in the nearby venous valves of the jugular and subclavian veins (**Figure 1A**, arrowheads).

**Figure 1:**
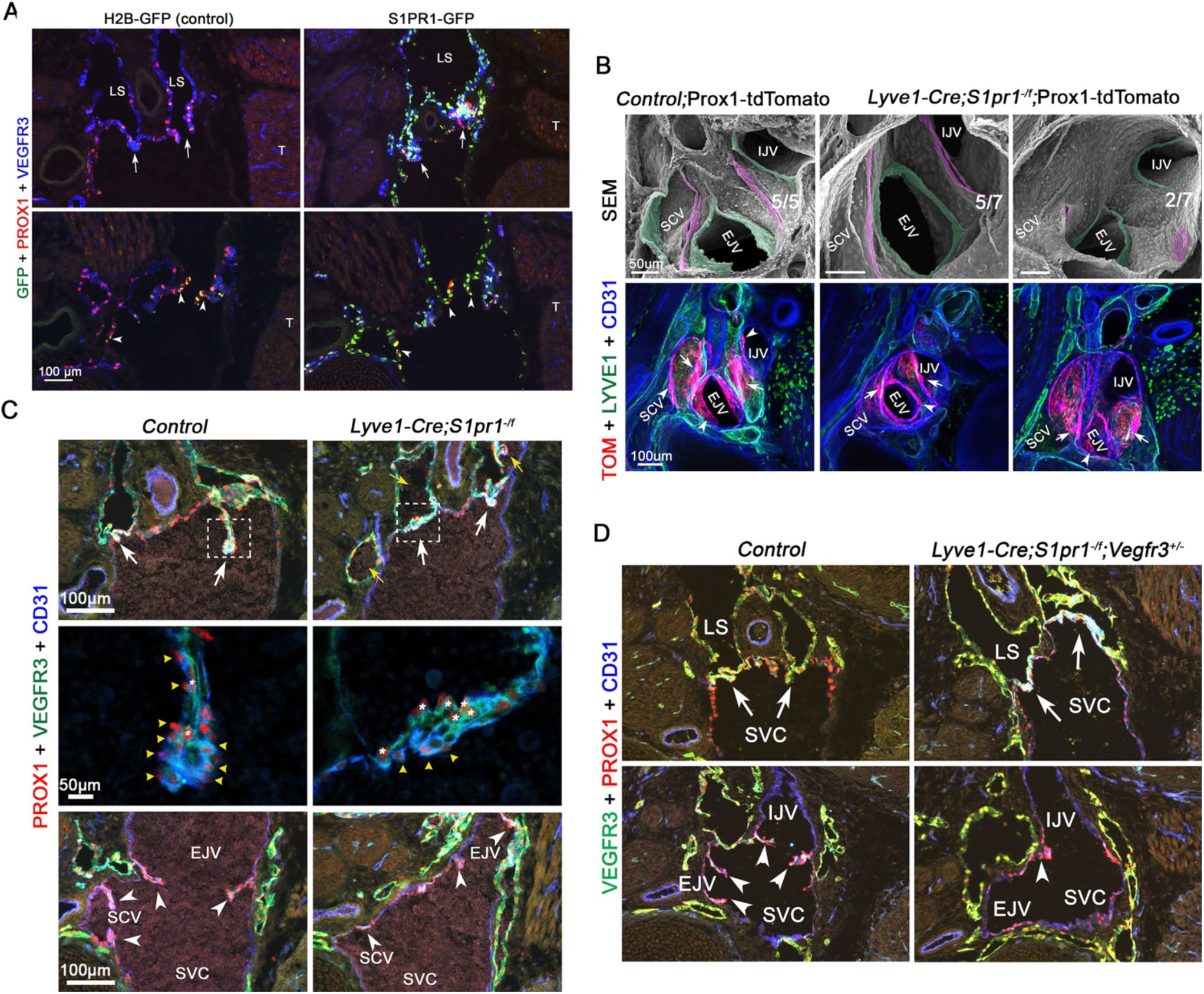
S1PR1 signaling is active in and is necessary for LVV and venous valve morphogenesis. (A) E16.5 S1PR1-GFP and H2B-GFP (control) littermates were frontally sectioned and analyzed. GFP^+^ cells were observed in the LVVs (arrows) and venous valves (arrowheads) of S1PR1-GFP embryos indicating that S1PR1 signaling is active in these structures. Only few GFP^+^ cells were observed in H2B-GFP embryos. (B) 500 μm thick transverse section of E16.5 Prox1-tdTomato and *Lyve1-Cre;S1pr1^-/f^*; Prox1-tdTomato embryos were prepared using a vibratome and whole mount immunohistochemistry and confocal imaging was performed to visualize the LVVs and venous valves (bottom row). Subsequently, the same samples were processed and analyzed by SEM (top row). Normal-looking LVVs (arrows or pseudo colored in magenta) and venous valves (arrowheads or pseudo colored in green) were seen in control embryos. Mutant embryos appeared to have substantially fewer valvular endothelial cells. When present the mutant cells appeared disorganized and arrested at the periphery of the vessels. (C, D) E16.5 control, *Lyve1-Cre;S1pr1^-/f^*and *Lyve1-Cre;S1pr1^-/f^*;*Vegfr3^+/-^* embryos were frontally sectioned (12 μm thick) and analyzed by IHC. LVVs and venous valves of control embryos had invaginated into the veins. Higher magnification images of control LVVs (boxed areas and the panels below in C) revealed that VEGFR3^hi^;Prox1^hi^ lymphatic endothelial cells (asterisks) were located in between two layers of VEGFR3^Low^;Prox1^hi^ valvular endothelial cells (yellow arrowheads). In contrast, invagination was substantially reduced in the LVVs and venous valves of mutant embryos and the organization of the two cell types was defective. Blood cells were observed within the lymph sacs of mutant embryos (yellow arrows in C). (D) The invagination defect of *Lyve1-Cre;S1pr1^-/f^* embryos was not rescued by *Vegfr3*-heterozygosity. Statistics: (A, C, D) n=5 embryos per genotype; (B) n=5 controls and n=7 *Lyve1-Cre;S1pr1^-/f^* LVV complexes. Abbreviations: LS, lymph sacs; T, thymus; EJV, external jugular vein; SVC, superior vena cava; SCV, subclavian vein; IJV, internal jugular vein

We used *Lyve1-Cre* to delete *S1pr1* from embryos (22, 39). *Lyve1-Cre* is active in the jugular vein, and the developing LECs (29, 35). Fluorescent/SEM correlative microscopy revealed that the LVV and venous valve leaflets of E16.5 *Lyve1-Cre; S1pr1^-/f^* embryos were either absent or much shorter than those of the control valves (**Figure 1B**). IHC performed on cryosections further revealed that the valvular endothelial cells were specified but failed to organize and invaginate into the veins (**Figure 1C**). Additionally, fewer LVs were observed in the dermal lymphatic vessels of *Lyve1-Cre; S1pr1^-/f^* embryos when compared to control littermates (**Supplementary** Figure 1). Thus, S1PR1 is necessary for the development of LVVs, LVs and venous valves.

We previously showed that the deletion of *S1pr1* from LECs resulted in excessive lymphatic vessel branching in embryonic skin, and that this phenotype can be ameliorated by deleting one allele of *Vegfr3* (39). To determine the effect on dermal LV defects in S1PR1-mutants we analyzed E16.5 *Lyve1-Cre; S1pr1^-/f^*; *Vegfr3^+/-^* embryos and determined that the dermal LV defects of S1PR1-mutants were not rescued by *Vegfr3*-heterozygosity (**Supplementary** Figure 1). The LVV and venous valve defects of S1PR1-mutants were also not rescued by *Vegfr3*-heterozygosity (**Figure 1D**). Together these data revealed that S1PR1 regulates the development of LVs, LVVs and venous valves in a *Vegfr3*-independent manner.

### S1PR1 regulates the postnatal development of lymphatic valves and maintains the functioning of lymphatic valves in the ileum-draining lymphatic vessels

*Lyve1-Cre; S1pr1^-/f^* mice have mispatterned mesenteric blood and lymphatic vessels due to Cre activity in both cell types in this organ (39). Therefore, we used alternative approaches to investigate the role of S1PR1 in postnatal LV development. First, we used the S1PR1-GS mice to test if S1PR1 signaling is active in the mesenteric LVs. Analysis of the mesenteries of P10 S1PR1-GS pups revealed that S1PR1 signaling is active in the mesenteric collecting lymphatic vessels although it is most potently expressed in LVs (**Figure 2A**, arrowheads). Next, to prevent mesenteric blood vascular defects we used transgenic Prox1-CreERT2 [Tg(Prox1-CreERT2)] to delete *S1pr1* from LECs (30, 41). To delete *S1pr1* we fed tamoxifen (TM) to Tg(Prox1-CreERT2);*S1pr1^f/f^* (*S1pr1^iΔLEC^*) pups and their control littermates. Postnatal day (P)1 pups were fed 1 μl of 20 mg/ml TM, P3 pups 3 μl and so on until P7. We generated P10 *S1pr1^iΔLEC^*(TM@P1-7) pups and confirmed the downregulation of S1PR1 in the mesenteric lymphatic vessels (**Supplementary** Figure 2A). A significant reduction in the number of mesenteric LVs was observed in *S1pr1^iΔLEC^* pups (**Figure 2B**). Furthermore, LVs were more severely reduced in the mesenteric lymphatic vessels that drain the distal small intestine (ileum) when compared to the proximal small intestine (duodenum and jejunum). The valve markers PROX1, FOXC2, GATA2 and integrin-α9 were normally expressed in the remaining LVs (**Supplementary** Figure 2A-C), although the expression of the gap junction molecule connexin-37 (CX37) appeared to be reduced (**Figure 2C**). Thus, S1PR1 is necessary for the postnatal development of mesenteric LVs.

**Figure 2:**
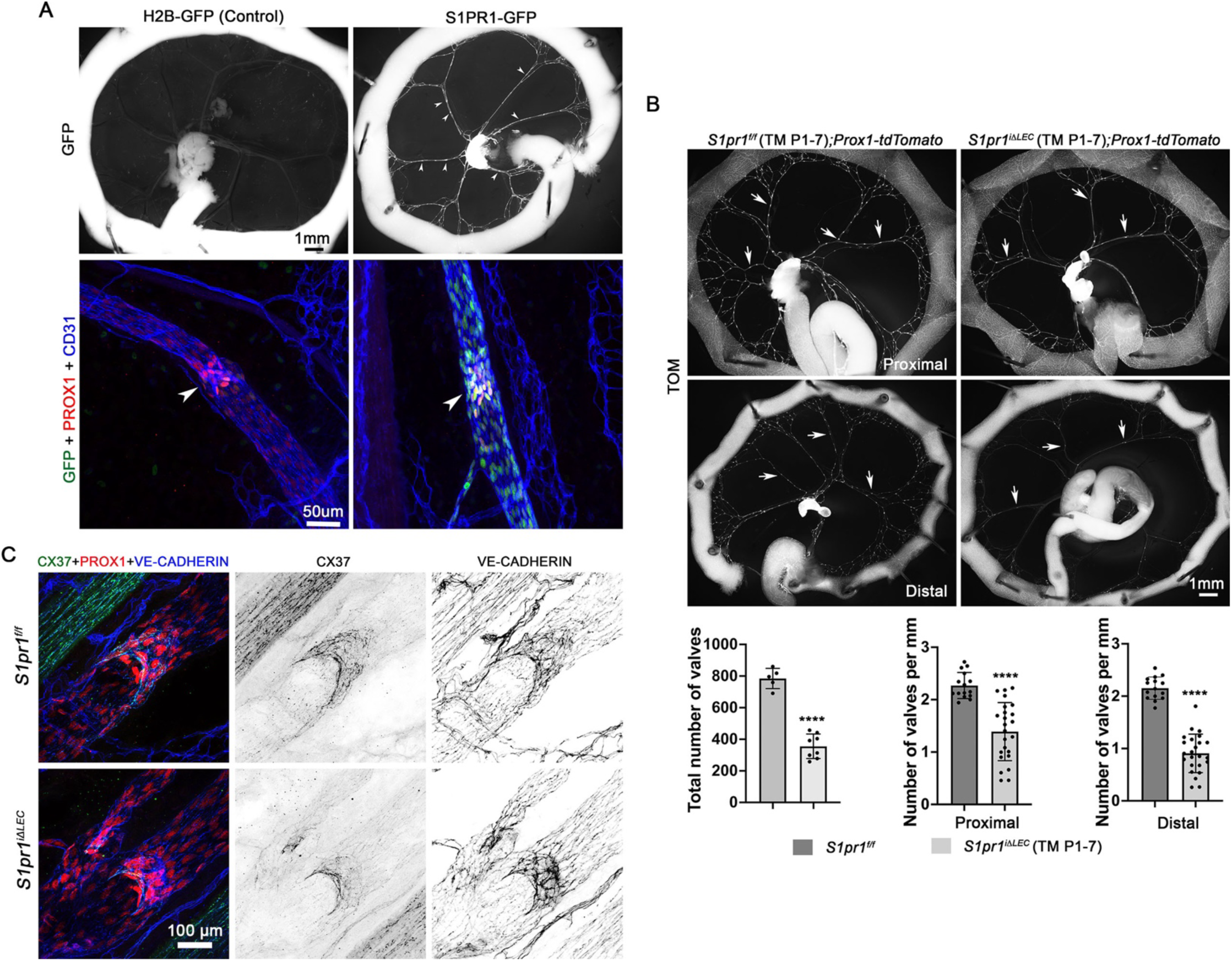
S1PR1 signaling is active in LVs and is necessary for LV development. (A) The mesenteries of P10 S1PR1-GFP and H2B-GFP littermates were analyzed. Collecting lymphatic vessels of S1PR1-GFP pups were GFP^+^ and GFP expression was stronger in the LVs (arrowheads). GFP expression was not observed in H2B-GFP pups. (B) The mesenteries of P10 *S1pr1^f/f^*;Prox1-tdTomato and *S1pr1^iΔLEC^*;Prox1-tdTomato pups that were administered tamoxifen from P1-7 were analyzed. Mesenteric tissue that was connected to duodenum and jejunum was considered “anterior” and that which is connected to ileum and cecum was considered as “posterior”. LVs (puncta) in the anterior, posterior and the entire gut were counted and quantified. LVs were significantly reduced in *S1pr1^iΔLEC^*;Prox1-tdTomato pups. The reduction appeared to be more severe in the posterior section of the gut. (C) The mesenteric lymphatic vessels and LVs of P10 pups that were administered tamoxifen from P1-7 were analyzed using the indicated antibodies. Connexin-37 (CX37) expression appeared to be reduced in the remaining LVs of *S1pr1^iΔLEC^* pups. Statistics: (A) n=9 S1PR1GFP and n=3 H2B-GFP pups; (B) n=5 control and n=8 *S1pr1^iΔLEC^* pups. Three anterior vessels and three posterior vessels from each mesentery were analyzed to quantify the valve density. Each dot represents a vessel on the graph; (C) n=3 per genotype. The graphs were plotted as Mean ± SD. Unpaired t test was performed for the statistical analysis. **** p<0.001.

To determine whether LVs are defective in other lymphatic vascular beds, we analyzed the ears of 3-month-old *S1pr1^iΔLEC^* (TM@P1-7) mice and determined that the dermal lymphatic vessels had a significantly higher number of branch points, but fewer number of LVs when compared to their control littermates (**Figure 3A**). Thus, S1PR1 is necessary for the development of dermal LVs. To determine if S1PR1 is necessary for the maintenance of already formed LVs we administered TM by gavage to 8-weeks-old mice for 3 consecutive days. The *S1pr1^iΔLEC^* (TM@8w) mice and their control littermates were analyzed 4 weeks later. No obvious increase in lymphatic vessel branch point density or reduction in LV numbers was observed in the ear lymphatic vessels (**Figure 3B**). Therefore, S1PR1 is not necessary to maintain already formed dermal LVs.

**Figure 3:**
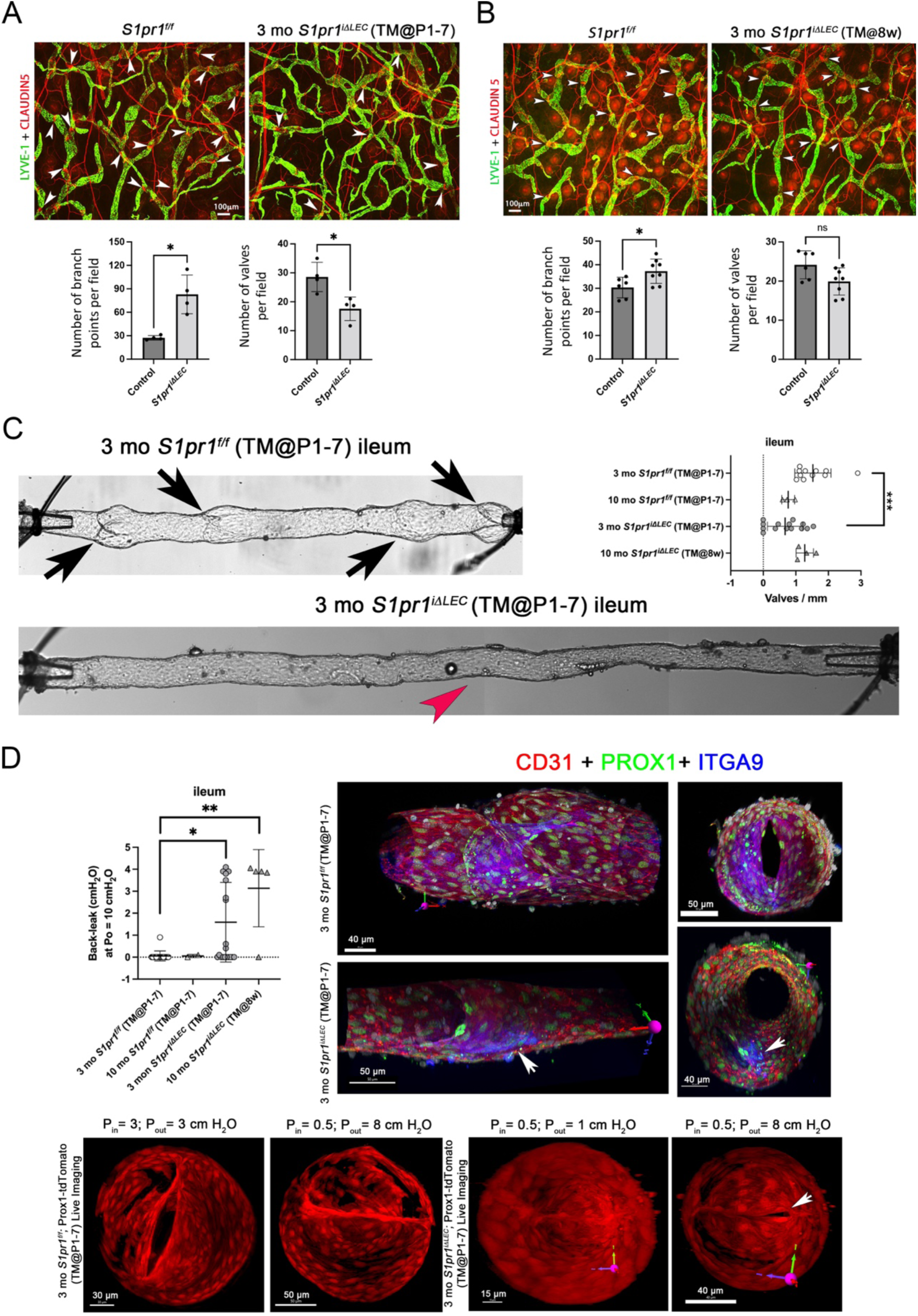
LVs are reduced in numbers and defective in *S1pr1^iΔLEC^* mice. (A, B) 3-month-old *S1pr1^iΔLEC^* mice that were treated with tamoxifen either from P1-P7 (A) or at 8 weeks of age (B) were studied. (A) Dermal lymphatic vessels in the ears of *S1pr1^iΔLEC^* (TM@P1-P7) mice had more branches and fewer Claudin-5^hi^ LVs (arrowheads) when compared to control littermates. (B) *S1pr1^iΔLEC^* (TM@8w) mice had elevated number of branch points, but did not have any obvious reduction in LVs. (C) Representative image of a LV (arrow) in the terminal-ileum draining mesenteric lymphatic vessel of a control mouse is shown. A corresponding lymphatic vessel from a *S1pr1^iΔLEC^* (TM@P1-7) mouse lacking LVs is also shown (red arrowhead indicates where small nubs remain from a valve). The graph shows that the LV density is significantly reduced in *S1pr1^iΔLEC^* (TM@P1-7), but not *S1pr1^iΔLEC^* (TM@8w) mice. (D) Ex-vivo analysis of LVs in the ileum draining lymphatic vessels. LVs of control, *S1pr1^iΔLEC^* (TM@P1-7) and *S1pr1^iΔLEC^* (TM@8w) mice were analyzed for back leak. The graph shows that the LVs in the terminal-ileum draining lymphatic vessels of mutant mice were significantly leaky irrespective of the time of gene deletion. Following ex-vivo analysis isolated vessel IHC was performed for the indicated markers, imaged by confocal microscopy and 3D reconstructed. LVs with 2 symmetrical leaflets were observed in control mice. Only one leaflet was observed in the leaky valve from this mutant. Control and “high-pressure leak” LVs with Prox1-tdTomato reporter were imaged live under various input (Pin) and output (Pout) pressures. LVs remained open when input and output pressures were equal. Both control and mutant LVs closed when Pout was slightly elevated. The control LV remained closed when Pout was increased to 8 cm H_2_O. In contrast, a gap remained in the mutant LV, resulting in back leak. Statistics: (A, B) Each dot represents an individual mouse. The graphs were plotted as Mean ± SD. Unpaired t test was performed for the statistical analysis. * p<0.05. (C) LV density was measured in ileum-draining lymphatic vessels harvested from n=11 3-mo *S1pr1^f/f^* (TM@P1-P7), n=3 10-mo *S1pr1^f/f^* (TM@8w), n=14 3-mo *S1pr1^iΔLEC^* (TM @ P1-P7) and n=4 10-mo *S1pr1^iΔLEC^* (TM@8w) mice. A one-way ANOVA withTukey’s post-hoc test was performed to determine significance. ***p<0.001. (D) Each dot represents an individual LV harvested from n=16 3-mo *S1pr1^f/f^* (TM@P1-P7), n=2 10-mo *S1pr1^f/f^* (TM@8w), n=21 3-mo *S1pr1^iΔLEC^* (TM @ P1-P7) and n=5 10-mo *S1pr1^iΔLEC^* (TM@8w) mice. A non-parametric Kruskal-Wallis test with Dunn’s post-hoc test was performed to determine significance. ** p<0.01. Live imaging and whole-mount IHC were performed using n=2 3-mo *S1pr1^f/f^*;Prox1-tdTomato (TM@P1-P7),and n=3 3-mo *S1pr1^iΔLEC^*;Prox1-tdTomato (TM @ P1-P7) LVs.

We analyzed the mesenteric lymphatic vessels of 3-month-old control and *S1pr1^iΔLEC^* (TM@P1-7) mice. The ileum-draining lymphatic vessels from control mice had 1-3 LVs per mm of vessel (**Figure 3C**). In contrast, lymphatic vessels from *S1pr1^iΔLEC^*(TM@P1-7) mice had fewer LVs and some were devoid of LVs (**Figure 3C**, red arrowhead). However, no obvious reduction in LV numbers was observed in 10-month-old *S1pr1^iΔLEC^* (TM@8w) indicating that S1PR1 is necessary for the formation, but not maintenance of LVs.

Next, mesenteric lymphatic vessels from various regions of the gut were isolated and cannulated to test LV function as described previously (42). LVs in the duodenum-, jejunum- or ileum-draining lymphatic vessels of 3- and 10-month-old control mice closed upon elevation of outflow pressure (Pout) and completely prevented back leak (**Figure 3D** and **Supplementary** Figure 3A**, B)**. Most LVs in the duodenum- and jejunum-draining lymphatic vessels of *S1pr1^iΔLEC^* (TM@P1-7) mice were also normal **(Supplementary** Figure 3A**, B)**. In contrast, significant numbers of LVs in the ileum-draining lymphatic vessels of *S1pr1^iΔLEC^* (TM@P1-7) mice were leaky (**Figure 3D**). We also analyzed the ileal LVs of 10-month-old *S1pr1^iΔLEC^*mice (TM@8w). Although these mice had normal numbers of LVs, they were significantly leaky (**Figure 3D**). Thus, S1PR1 is necessary to maintain the normal functioning of ileal LVs.

To determine the structural defects in leaky LVs we performed live imaging on LVs immediately after conducting valve tests. We harvested LVs from Prox1-tdTomato and *S1pr1^iΔLEC^*; Prox1-tdTomato (TM@P1-7) mice and imaged them live at various levels of inflow pressure (Pin) and Pout. Control LVs closed at low Pout and remained tightly closed at high Pout (**Figure 3D**). In contrast, a mutant LV closed at low Pout but developed gaps at high Pout (**Figure 3D**, arrow). We also performed IHC on isolated vessels after fixation and imaged them by confocal microscopy followed by 3D-reconstruction of the LVs. While control valves had 2 symmetrical leaflets, a leaky valve from a *S1pr1^iΔLEC^*(TM@P1-7) mouse had only one leaflet (**Figure 3D**, arrows). Another leaky valve appeared to have three leaflets (**Supplementary** Figure 4). One or both leaflets were abnormally elongated at their insertion points in the wall in two other leaky LVs (**Supplementary** Figure 4). These results indicated that S1PR1 is necessary to maintain the integrity of LVs. The significant heterogeneity in LV defects is consistent with the defects that were observed in LVVs (**Figure 1B**).

In summary, S1PR1 is necessary for the postnatal development of LVs, S1PR1 regulates LV morphogenesis that is necessary to prevent back-leak under adverse pressure gradient and LVs in the ileum are more sensitive to the loss of S1PR1 when compared to the LVs in the proximal mesentery.

### Lymphatic drainage is defective and tertiary lymphoid organs are present in the mesenteries of S1pr1^iΔLEC^ mice

Recently, TLOs were found to form in the vicinity of LVs in the terminal ileum of *Tnf^+/11ARE^* mice, a model for ileitis (11, 26). TNFα inhibited the expression of numerous valve regulatory molecules in HLECs cultured under static or OSS condition (11). Therefore, it was proposed that TNFα inhibits LV function and lymphatic drainage resulting in the accumulation of immune cells in the mesentery and subsequently TLO formation. As the LVs were defective in the terminal ileum of *S1pr1^iΔLEC^* mice we wanted to determine if lymphatic drainage is defective in these mice and if they develop TLOs.

We injected FITC-conjugated dextran (molecular weight = 2M Kd) into the muscle layer of the gut wall and/or the Peyer’s patches of anesthetized 10-month-old mice and performed live imaging to visualize the flow of fluorescent dye. In control mice, the dye was drained by the mesenteric collecting lymphatic vessels quickly and in a unidirectional manner (**Figure 4**, arrows and **Supplementary Movie 1**). In contrast, in *S1pr1^iΔLEC^* mice dye drained into numerous nodules that appeared to slow down the flow (**Figure 4**, arrowheads and **Supplementary Movie 2, 3**). Additionally, dye often appeared to flow in the retrograde direction from nodule to nodule (**Figure 4**, yellow arrows). However, no obvious leakage of dye was observed from the lymphatic vessels or nodules.

**Figure 4:**
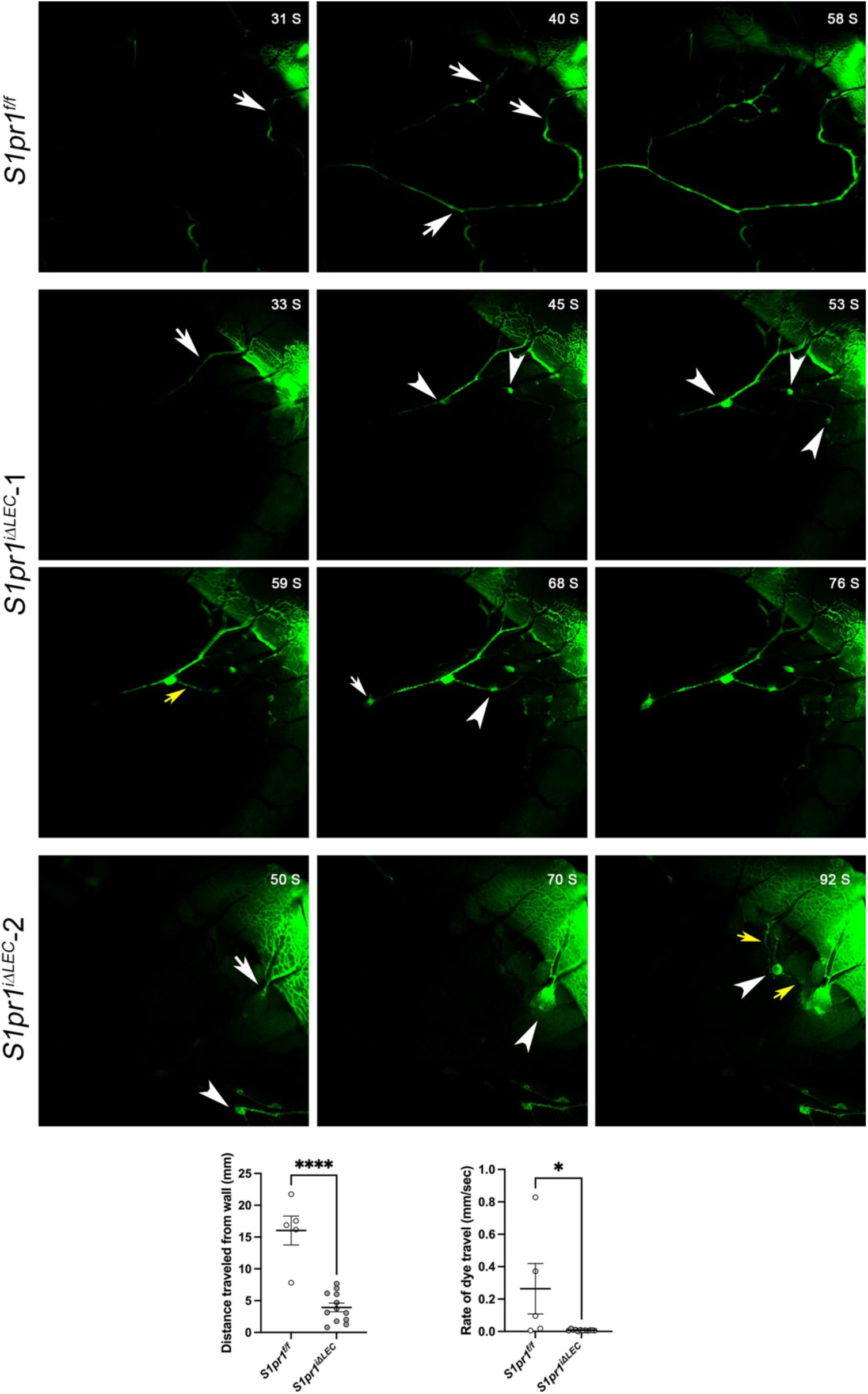
Lymphatic drainage is defective and obstructed by nodules in *S1pr1^iΔLEC^* mice. FITC-conjugated dextran (2M Kd) was injected into the muscle layer of the ileum and/or the Peyer’s patches of anesthetized 10-month-old *S1pr1^f/f^* and *S1pr1^iΔLEC^* mice that were treated with tamoxifen from P1-7. The flow of fluorescent dye through the mesenteric lymphatic vessels was visualized by live imaging. Time in seconds after injection is indicated on the top right corner of the panels. The dye rapidly drains through the lymphatic vessels in control mice (white arrows). In contrast, the dye accumulated in nodules that were connected to the lymphatic vessels (white arrowheads). Retrograde flow was also observed between the nodules (yellow arrows). The videos were analyzed to quantify the distance travelled by the dye and the rate of travel. The graphs show that these parameters were significantly reduced in *S1pr1^iΔLEC^* mice. Statistics: Images are representative of n=4 *S1pr1^f/f^* and n=5 *S1pr1^iΔLEC^* mice. Some samples were analyzed by injection at multiple sites. Each dot in the graph indicates an individual injection. Unpaired t tests were performed for the statistical analyses. * p<0.05; **** p<0.001.

The appearance and location of the nodules in the mesentery of *S1pr1^iΔLEC^*mice were reminiscent of TLOs that were reported in the terminal ileum of *Tnf^+/ΔARE^*mice (11, 26). Therefore, we characterized the mesenteric tissue by IHC with markers of immune, stromal and vascular cells. IHC for the LEC marker VEGFR3 revealed that the nodules were primarily located in the terminal ileum of *S1pr1^iΔLEC^* mice, as in *Tnf^+/ΔARE^*mice (**Figure 5A**). On average, 60-70 nodules that measured ∼300 μm in diameter were observed in *S1pr1^iΔLEC^*mice (**Figure 5B**). Some of the *S1pr1^iΔLEC^* mice had *R26^+/tdTomato^* reporter to permanently label PROX1^+^ lymphatic vessels at the time of tamoxifen injection. Expression of tdTomato, LYVE1, VEGFR3 and PROX1 revealed that the lymphatic vessels were wrapped around the nodules (**Figure 5C**). Furthermore, the *R26^+/tdTomato^* lineage-tracer revealed that these lymphatic vessels had originated from lymphatic vessels that existed at least as early as P1-7, the time of tamoxifen administration.

**Figure 5:**
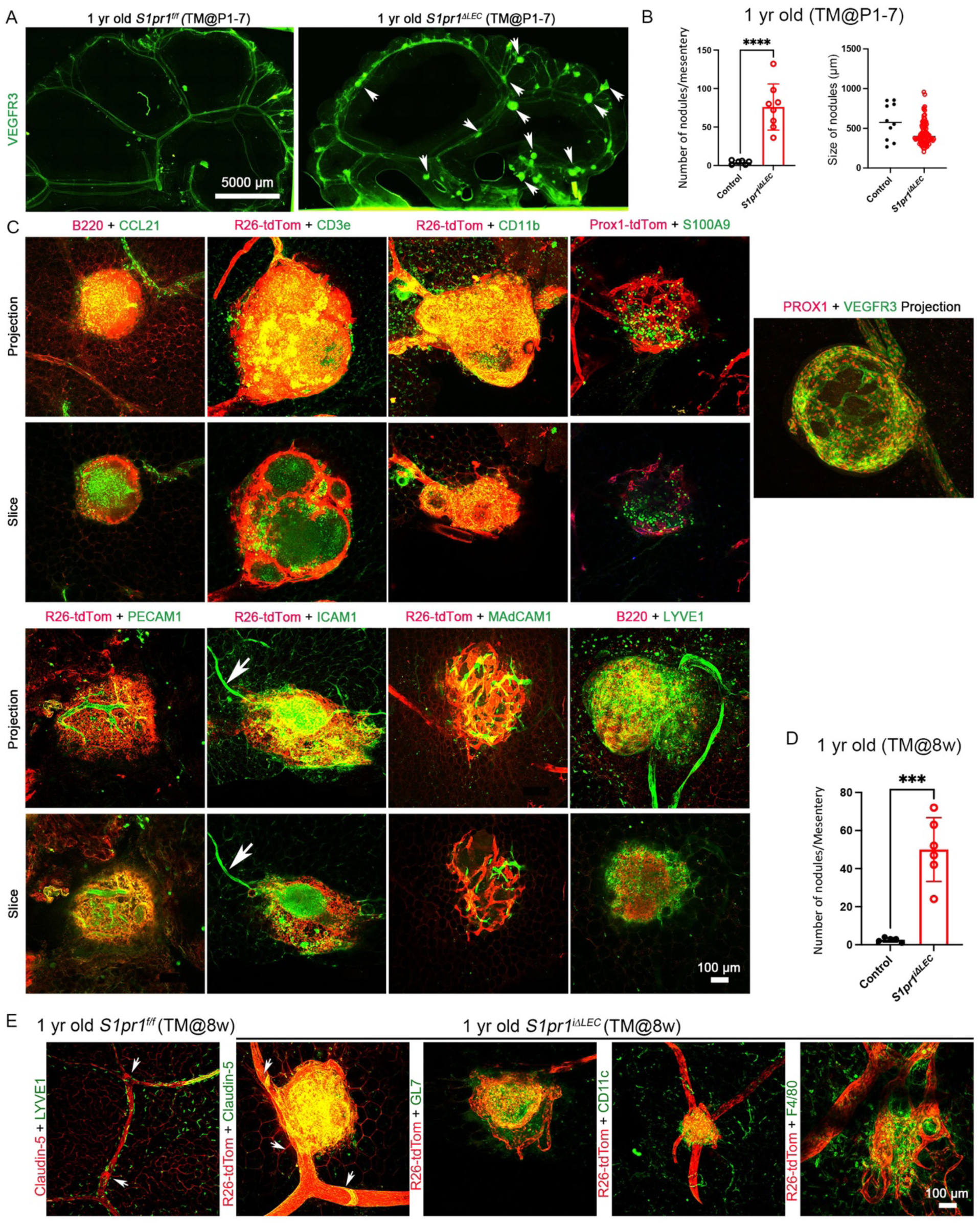
Tertiary lymphoid organs are present in the terminal ileum of *S1pr1^iΔLEC^* mice, and they are connected to the lymphatic vessels. (A) Mesenteric tissue along the ileum was harvested and analyzed using the lymphatic vessel marker VEGFR3. Large number of nodules were observed in *S1pr1^iΔLEC^* (TM@P1-7) mice, but not in control littermates. (B) The number and size of LYVE1^+^B220^+^ nodules were measured and quantified. (C) Mesenteries of 8-12-month-old control mice and *S1pr1^iΔLEC^* littermates that were treated with tamoxifen from P1-7 were analyzed using markers for the immune cells, stromal cells and endothelial cells. Some *S1pr1^iΔLEC^*mice had *R26^+/tdTomato^* reporter, the expression of which was permanently induced in the lymphatic vessels by tamoxifen activated CreERT2. Lymphatic vessels were labelled by VEGFR3, PROX1, LYVE1 and tdTomato. LYVE1 was also expressed in a subset of macrophages. B220, CD3e, CD11b and S100A9 are markers of B cells, T cells and myeloid lineage cells, and neutrophils respectively. CCL21 is a marker for lymphatic vessels and the fibroblast reticular cells within TLOs. PECAM1 labels all endothelial cells, but its expression is stronger in blood endothelial cells when compared to lymphatic endothelial cells. ICAM1 is expressed in HEVs, inflamed blood endothelial cells and a variety of immune cells. MAdCAM1 is a marker of HEVs. (D) TLO’s in the terminal ileum of 1 year old *S1pr1^f/f^* and *S1pr1^iΔLEC^* (TM@8W) mice were counted and plotted. (E) Mesenteric lymphatic vessels of *S1pr1^f/f^* mice had clear Claudin-5^+^ LVs (arrows) and weak LYVE1 expression. LVs were also observed in *S1pr1^iΔLEC^* (TM@8W) mice (arrows), although those that were close to the TLOs appeared defective. The TLOs had GL7^+^ germinal center B cells, CD11c^+^ dendritic cells, and F4/80^+^ macrophages. Statistics: (A) n=3 *S1pr1^f/f^* and n=3 *S1pr1^iΔLEC^* (TM@P1-7) mice; (B) n=6 *S1pr1^f/f^* and n=8 *S1pr1^iΔLEC^* mice. The size of the nodules was quantified, and each dot represents a nodule on the graph; (C, E) Representative images from 3-5 mice/genotype/marker; (D) n=6 *S1pr1^f/f^* and n=7 *S1pr1^iΔLEC^*(TM@8W) mice. Graphs were plotted as mean ± SD. Unpaired t test was performed for the statistical analysis. * p<0.05; **** p<0.001.

The nodules contained B220^+^ B cells, CD3e^+^ T cells, CD11b^+^ leukocytes, S100A9^+^ neutrophils, CCL21^+^ LECs and CCL21^+^ fibroblastic reticular cells (FRCs) that were located at the core of the nodules (**Figure 5C**). IHC for the pan-endothelial marker CD31/PECAM1 revealed the presence of tdTomato^-^ blood vessels within TLOs (**Figure 5C**). Lymphocytes enter the LNs from the blood circulation via HEVs (3); some of the tdTomato^-^ blood vessels expressed the HEV marker MAdCAM1 (**Figure 5C**). Leukocytes extravasate from blood vessels through the interaction of CD11b with adhesion molecules such as ICAM1 that are expressed on inflamed endothelial cells (43) and HEVs (44). ICAM1 is also expressed in LTo cells and some leukocytes (19). ICAM1 was identified in blood vessels both within and outside the nodules of *S1pr1^iΔLEC^* mice (**Figure 5C, arrow**). On the other hand, ICAM1 expression was patchy in the tdTomato^+^ lymphatic vessels. Based on the expression pattern of various markers we concluded that the nodules in the terminal ileum of *S1pr1^iΔLEC^*mice were TLOs that contain lymphatic vessels, inflamed blood vessels, HEVs, B cells, T cells, myeloid cells, and FRCs.

Finally, we harvested the mesenteries of 1-year-old *S1pr1^iΔLEC^*mice that were exposed to tamoxifen at 8-weeks of age and determined that they too had TLOs with CD11c^+^ DCs, GL7^+^ germinal center B cells and F4/80^+^ macrophages in the terminal ileum (**Figure 5D, E**). Thus, S1PR1 is constantly required to prevent TLO formation in the terminal ileum and corelates with the presence of defective LVs.

### LV development and TLO formation are regulated by cell-autonomous S1PR1 signaling in LECs

S1P is the ligand for S1PR1, and it is generated by a complex metabolic pathway consisting of numerous intermediates and regulatory enzymes (45). The final step of S1P synthesis is mediated by sphingosine kinases 1 and 2 (SPHK1/2), which convert sphingosine to S1P. S1P is secreted from hematopoietic cells by the transporter MFSD2B and from endothelial cells by the transporter SPNS2 (37). Deletion of *Sphk1/2* or *Spns2* from LECs downregulates S1P levels in lymph (22, 46). Thus, LECs are the primary source of S1P in lymph.

We generated P10 *Lyve1-Cre;Sphk1^-/f^;Sphk2^-/-^*(*Sphk1/2 ^ΔLEC^*) pups and analyzed their mesenteric LVs. *Sphk1/2 ^ΔLEC^* pups were phenotypically similar to *S1pr1^iΔLEC^*pups and had significantly fewer mesenteric LVs (**Figure 6A**). Analysis of the ears of 3-month-old *Sphk1/2 ^ΔLEC^* mice revealed fewer dermal LVs (**Figure 6B**). Additionally, the dermal lymphatic vascular density was increased in *Sphk1/2 ^ΔLEC^* mice. These data confirmed our hypothesis and showed that LV development is regulated by S1P produced by LECs, which could be activating S1PR1 signaling in an autocrine or paracrine manner. LVVs and venous valves of *Sphk1/2 ^ΔLEC^* mice were not analyzed as these valves are exposed to S1P derived from blood endothelial cells and hematopoietic cells.

**Figure 6:**
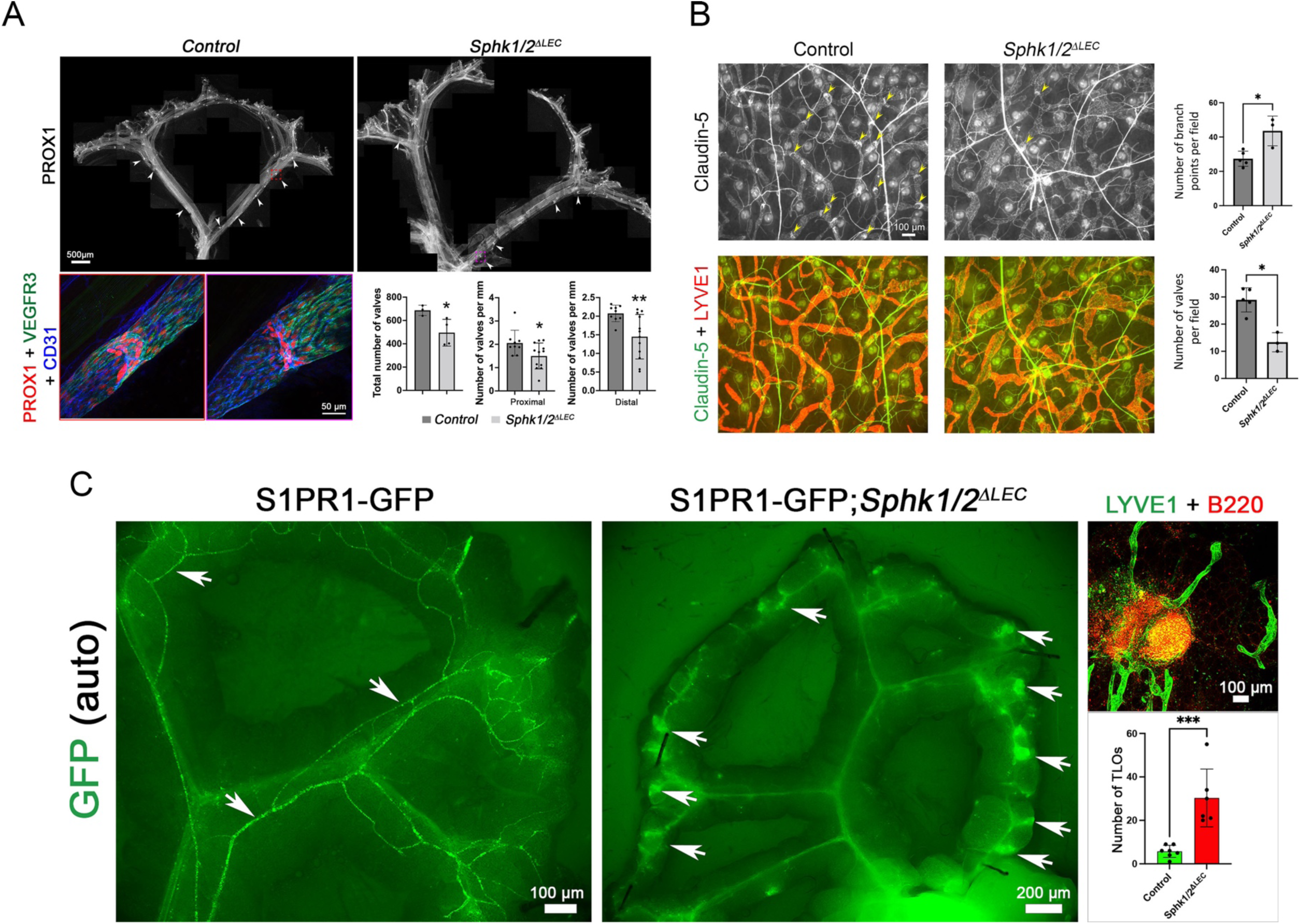
Autocrine or paracrine S1PR1 signaling regulates LV development and TLO formation. (A, B) Mesenteric (A) and dermal (B) lymphatic vessels of *Sphk1/2^ΔLEC^* mice, in which S1P synthesis in lymphatic endothelial cells was ablated, were analyzed. (A) Mesenteric lymphatic vessels of P10 *Sphk1/2^ΔLEC^* pups had fewer LVs and those that were remaining appeared immature. Representative valves are within dotted boxes and their enlarged images are shown below. (B) Dermal lymphatic vessels of 3-month-old *Sphk1/2^ΔLEC^* mice had fewer LVs (yellow arrows) and more branches per field. (C) The mesenteries of 1-year-old S1PR1-GFP and S1PR1-GFP;*Sphk1/2^ΔLEC^* mice were analyzed. GFP autofluorescence was observed in the lymphatic vessels of S1PR1-GFP mice (arrows). Weaker GFP expression was observed in blood vessels. In contrast, lymphatic vessels could not be identified based on GFP autofluorescence in S1PR1-GFP;*Sphk1/2^ΔLEC^* mice. However, GFP expression was observed in blood vessels and in nodule-like structures (arrows). IHC for LYVE1 and B220 revealed that the GFP^+^ nodules were TLOs that contain lymphatic vessels and B cells. Statistics: (A) n=3 control and n=4 *Sphk1/2^ΔLEC^* pups. Three proximal vessels and three distal vessels from each mesentery were analyzed to quantify the valve density. Each dot in the graph represents a vessel; (B) n=4 controls and n=3 *Sphk1/2^ΔLEC^* mice. Three fields were imaged from one of the immunostained ears of the mice. All 3 fields were used to quantify the number of branch points and vessel diameter. One of the fields was used for counting the number of LVs. (C) n=7 S1PR1-GFP and n=8 *Sphk1/2^ΔLEC^*;S1PR1-GFP mice. Graphs were plotted as mean ± SD. Unpaired t test was performed for the statistical analysis. * p<0.05; ** p<0.01; *** p<0.005; **** p<0.001.

To determine if LEC-derived S1P inhibits TLO formation we analyzed the mesenteries of 1 year old S1PR1-GS and S1PR1-GS*; Sphk1/2 ^ΔLEC^* mice. The lymphatic vessels of control mice were GFP^+^, demonstrating that S1PR1 signaling is active in LECs (**Figure 6C**, arrows). In contrast, GFP expression was downregulated in the lymphatic vessels of *Sphk1/2 ^ΔLEC^*mice (**Figure 6C**) as previously demonstrated (47, 48). Additionally, numerous GFP^+^ clusters were observed in the terminal ileum of *Sphk1/2 ^ΔLEC^* mice (**Figure 6C**, arrows). IHC for B220 and LYVE1 revealed these clusters to be TLOs (**Figure 6C**). These data indicate that cell autonomous S1PR1 signaling in LECs inhibits TLO formation in the terminal ileum.

In summary, autocrine or paracrine S1PR1 signaling in LECs regulates LV development and inhibits TLO formation in the terminal ileum. Importantly, TLOs formed at sites that lacked LVs or had defective LVs, thus suggesting that LV defects could contribute to TLO formation as hypothesized previously (11).

### Deletion of S1PR1 from the lymphatic vasculature does not result in epithelial dysplasia, epithelial inflammation or microbial dysbiosis

TLOs are observed in the mesenteries of Crohn’s disease patients and in a mouse model for ileitis (11, 12, 26). TLOs are thought to form due to chronic inflammation although this possibility has not been tested (1). Whether TLOs can promote or aggravate the disease by causing epithelial inflammation and tissue damage is also not known.

The body weight, number of Peyer’s patches, spleen size and mesenteric lymph node size of 1-year-old *S1pr1^iΔLEC^* mice were indistinguishable from those of control littermates (**Supplementary** Figure 5). H&E staining did not reveal differences in epithelial morphology of the terminal ileum of control and *S1pr1^iΔLEC^* mice (**Supplementary** Figure 6A), nor did IHC reveal any obvious increase in the infiltration of S100A9^+^ neutrophils or B220^+^ B cells in the ileum of mutant mice **(Supplementary** Figure 6B). Of a panel of cytokines measured in the serum of *S1pr1^iΔLEC^* mice, we only observed a modest increase in IL-7 **(Supplementary** Figure 6C). Although a trend to an increase was also observed in 11 other inflammatory cytokines including TNFα and IL-1β, these changes were not significant **(Supplementary** Figure 6C).

GWAS studies have implied that an abnormal immune response to commensal bacteria is the primary cause of Crohn’s disease (49–51). Crohn’s disease is associated with microbial dysbiosis, in which a reduction in beneficial organisms (e.g., *Faecalibacterium prausnitzii*, of the phyla Firmicutes) and an expansion of pathological microorganisms (e.g., *Escherichia coli, of* the phyla Proteobacteria) is observed (49, 52).

Microbiota dysbiosis was also observed in mouse models of ileitis (53–55). We performed shotgun metagenomic analysis of fecal pellets from the terminal ileum of 12-month-old *S1pr1^iΔLEC^* mice and littermate controls. No striking differences were observed in the bacterial contents of the mutants when compared to their control littermates (**Supplementary** Figure 6D). Thus, microbial dysbiosis is not observed in *S1pr1^iΔLEC^*mice.

In summary, *S1pr1^iΔLEC^* mice do not have the characteristics of ileitis. Thus, TLOs in the mesentery of *S1pr1^iΔLEC^*mice are a sign of subclinical local inflammation that does not result in systemic inflammation, epithelial damage or microbial dysbiosis.

### S1PR1 regulates cytoskeletal organization, oscillatory shear stress response, and the expression of valve-regulatory genes

Oscillatory shear stress (OSS) can enhance the expression of molecules such as the transcription factor FOXC2 and the gap junction molecule CX37, both of which are critical for lymphatic vessel maturation and LV development (31, 56–58). LECs require the ion channel PIEZO1 and the adherens junction molecule VE-Cadherin for sensing OSS and activating the expression of valve-regulatory molecules (36, 59). However, the mechanisms by which LECs sense and transduce OSS are not fully understood.

S1PR1 can regulate the cytoskeleton, stabilize adherens junctions and mediate shear stress responses in blood endothelial cells (60, 61). We previously showed that S1PR1 is necessary for cytoskeletal organization in primary human lymphatic endothelial cells (HLECs) (39). We also showed that S1PR1 can regulate the laminar shear stress response in HLECs (39). Hence, we hypothesized that S1PR1 signaling regulates the expression of valve-regulatory molecules in response to OSS.

We performed IHC for the expression of actin and VE-Cadherin in siControl- and siS1PR1-transfected HLECs grown under static or OSS conditions. As reported previously, control HLECs became more spherical in response to OSS and had thicker cortical actin fibers (31, 62). Cell-cell junctions changed from linear junctions under static conditions to overlapping junctions (**Figure 7A**). In contrast, siS1PR1-transfected HLECs were elongated in shape under both static and OSS conditions (**Figure 7A**). Stress fibers that crisscrossed the cytoplasm were the predominant type of actin that was observed. Furthermore, siS1PR1-transfected HLECs maintained linear cell-cell junctions despite exposure to OSS. These results demonstrated that S1PR1 is necessary for cytoskeletal architecture and adherens junction assembly in OSS-exposed HLECs.

**Figure 7:**
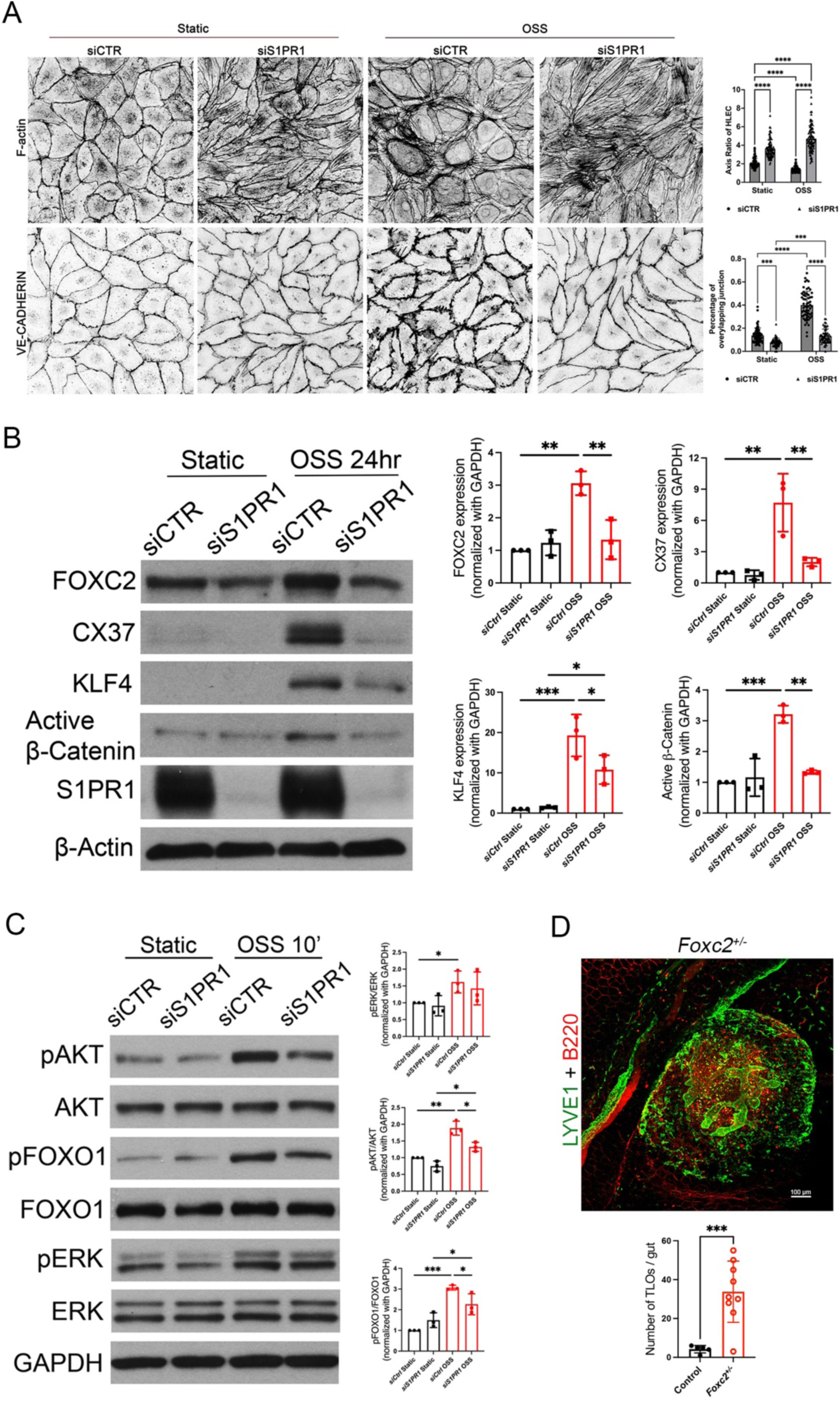
S1PR1 regulates OSS response and the expression of valve regulatory genes in HLECs. (A) HLECs were transfected with siControl or siS1PR1 and grown for 24 hrs under static condition for downregulating S1PR1. Subsequently, cells were cultured under static or OSS for 24 hrs. Cells were immunostained for F-actin or VE-Cadherin. F-actin was primarily located along the cell wall (cortical actin) of control cells under both static and OSS conditions. Control HLECs became more spherical, and cortical actin expression appeared to be increased by OSS. In contrast, siS1PR1 transfected HLECs appeared elongated and had increased expression of stress fibers. The percentage of VE-cadherin^+^ overlapping cell junctions was increased by OSS, and this enhancement was abolished by siS1PR1. (B) HLECs were cultured as described above and western blotting was performed for the indicated proteins. OSS induced the expression of the shear stress responsive transcription factor KLF4 and the valve-regulatory molecules active-β-catenin, FOXC2 and Connexin-37. Knockdown of S1PR1 significantly inhibited the expression of these molecules. (C) HLECs were transfected with siControl or siS1PR1 and grown for 48 hrs under static condition for downregulating S1PR1. Subsequently, cells were cultured under static or OSS for 10 minutes. Cell lysates were western blotted for the indicated antibodies and quantified. pAKT, pERK and pFOXO1 were upregulated by OSS. siS1PR1 significantly downregulated the expression of pAKT and pFOXO1. (D) The mesenteric tissues from 1-year-old control and *Foxc2^+/-^* mice were analyzed by IHC for the indicated markers to identify and quantify TLOs. A representative TLO from a *Foxc2^+/-^* mouse is shown. Significantly higher number of TLOs were observed in the *Foxc2^+/-^*mice. Statistics: (A) The axis was measured in 30 cells and junction was analyzed in 20 cells in a single field from each of the 3 experiments. Each dot represents one cell in the graphs; (B) The blot is representative of 3 independent experiments. The data from all experiments was used to prepare the graphs; (C) The blot is representative of 3 independent experiments. The data from all experiments was used to prepare the graphs; (D) Each dot in the graph indicates an individual mouse. n=5 controls, n=9 *Foxc2^+/-^* mice. The graphs are shown as mean ± SD. Two-way ANOVA with Tukey’s post-hoc test was performed to determine statistical significance. * p<0.05; ** p<0.01; *** p<0.005; **** p<0.001.

We transfected HLECs with control siRNA or siS1PR1 and cultured them for 24 hours under static or OSS conditions. Cells were lysed and western blotting was performed to quantify the expression of FOXC2 and CX37. OSS induced the expression of FOXC2 and CX37 in control HLECs as reported previously (31). Knock down of S1PR1 significantly downregulated the expression of FOXC2 and CX37 (**Figure 7B).** We previously showed that OSS enhances Wnt/β-catenin signaling, which is necessary for valve development and FOXC2 expression (35). Accordingly, OSS enhanced the expression of active-β-catenin (**Figure 7B**). In contrast, active-β-catenin was not upregulated in siS1PR1-transfected HLECs (**Figure 7B**). Thus, S1PR1 regulates Wnt/β-catenin signaling and the expression of valve regulatory molecules in response to OSS.

Acute OSS promotes AKT phosphorylation in HLECs in a VE-Cadherin-dependent manner (59). In turn, pAKT phosphorylates and inhibits the transcription factor FOXO1 to derepress FOXC2 (63). We knocked down S1PR1 in HLECs and cultured them under static or OSS conditions for 10 minutes. Western blotting revealed that OSS enhanced the phosphorylation of AKT and ERK in control HLECs (**Figure 7C**). Knockdown of S1PR1 significantly reduced the phosphorylation of AKT and FOXO1 in response to OSS (**Figure 7C**).

In summary, S1PR1 regulates cytoskeletal- and adherens junction-organization in HLECs. S1PR1 also regulates AKT phosphorylation and the expression of valve regulatory molecules in response to OSS.

### Foxc2-heterozygous mice develop TLOs in the mesentery

As reported above, S1PR1 regulates the expression of several genes that are necessary for LV development among which FOXC2 is a central player. LVs do not develop in mice lacking FOXC2 (64). Heterozygous loss-of-function mutations in *FOXC2* are associated with lymphedema-distichiasis syndrome (LDS) (65, 66). LDS patients have incompetent venous valves (67). Whether their LVs are also defective is not known due to the difficulties associated with imaging their activity. *Foxc2^+/-^* mice, used as a model of LDS, possess ∼50% fewer mesenteric LVs (63). Slight leakage was observed in the remaining valves (63). Approximately 50% of LDS patients develop “multiple small nodules” in the mesentery (65). Whether these nodules are TLOs is not known although an excessive number of lymph nodes were reported to develop throughout the body of *Foxc2^+/-^* mice (68). Hence, we analyzed the mesenteries of *Foxc2^+/-^*mice and determined that they indeed have TLOs in the terminal ileum (**Figure 7D**). These data indicate that FOXC2 is a physiologically relevant target of S1PR1 during both LV development and TLO formation and suggest that LV defects could lead to TLO formation.

## DISCUSSION

We report several new discoveries in this work: 1. S1PR1 signaling is a novel regulator of LV development; 2. Differences exist between mesenteric LVs along the proximal to distal axis; 3. TLOs can form in the absence of severe inflammation or tissue damage, and 4. LV defects can result in TLO formation. A schematic working model based on our findings is presented in **Supplementary** Figure 7.

### S1PR1 and LV development

Existing data supports the role of OSS in valve development (31). OSS is sensed and transduced by the mechanosensory ion channel PIEZO1 (36). The endothelial adherens junction molecule VE-Cadherin is necessary for activating Wnt/β-catenin signaling and AKT phosphorylation in response to OSS (59). In turn, β-catenin forms a complex with PROX1 to promote FOXC2 expression, and pAKT promotes FOXC2 expression by inhibiting FOXO1 (63, 69, 70). In this work we have identified S1PR1 as a molecule that is necessary for the phosphorylation of AKT and FOXO1 in response to OSS and for the expression of the valve-regulatory molecules active β-catenin, FOXC2 and connexin-37. S1PR1 likely regulates the OSS response through the cytoskeletal architecture and adherens junction complex. Additional studies are necessary to delineate the relationship between S1PR1 and the other mechanosensory molecules such as PIEZO1 and VE-Cadherin. Our work also suggests that S1P is necessary for LV development and that it activates S1PR1 signaling in an autocrine or paracrine manner.

### Heterogeneity of mesenteric LVs

Previous publications have only evaluated LV formation and function in the duodenum or jejunum (42, 62, 63, 71, 72). We have identified some intriguing peculiarities in the LVs that drain the ileum when compared to those that are in the proximal sections of the GI tract. Neonatal deletion of S1PR1 results in a reduced number of mesenteric valves, especially in the ileal mesentery. Although LVs were reduced in number also in the proximal GI sections of the mutant mice, the remaining LVs did not degenerate in adulthood. However, LVs in the terminal ileum of mutant mice are functionally defective. This defect occurs even if *S1pr1* is deleted from adult mice. We currently do not know the reason for these regional differences in LV development and function. We speculate (below) that the microbial antigens and cytokines that are generated in response to these antigens could compromise LV function in the terminal ileum. S1PR1 signaling appears to be necessary to minimize the impact of these damaging factors.

### Mechanisms of TLO formation

TLOs were not observed in the skin or proximal GI tract of *S1pr1^iΔLEC^*mice. The ileo-cecal junction of the GI tract, where the TLOs were predominantly observed, has the highest density of microbiota in the foregut (73). Relevant to our work, dendritic cells engulf infiltrating bacteria, migrate from the gut to the mesenteric lymph nodes and Peyer’s patches via lymphatic vessels, and activate T cells and IgA-secreting B cells (49, 74). These activated lymphocytes exit the lymph nodes through lymphatic vessels, enter the blood stream, and return to the gut to restrict pathological organisms and maintain microbial homeostasis. We speculate that due to LV defects *S1pr1^iΔLEC^*, *Sphk1/2 ^ΔLEC^* and *Foxc2^+/-^* mice are unable to build a systemic immune response against microbiota in the mesenteric lymph nodes (**Supplementary** Figure 7). Consequently, TLO’s develop in the terminal ileum to respond against microbial antigens locally and prevent microbial dysbiosis.

### Limitations of the study

We speculate that S1PR1 inhibits TLO formation by regulating LV development and function and thus promoting efficient lymphatic drainage. In support of this possibility TLOs were found in *Foxc2^+/-^* mice that have LV defects. Additionally, TNFα can inhibit the expression of valve-regulatory molecules and TLOs frequently form near the LVs of *Tnf^+/11ARE^* mice (11). We predict that other mouse models and humans with compromised lymphatic drainage in the ileum due to LV or lymphatic vessel defects will also have TLOs in this location. However, we cannot rule out alternative reasons for TLO formation. For example, S1PR1 signaling can antagonize TNFα-induced inflammation in blood endothelial cells (75). S1PR1 can also inhibit the expression of pro-inflammatory molecules such as *Irf8, Lbp, Il7, Il33, Ccl21* and *Tnfaip8l1* in LECs of mice (48). S1PR1 inhibits the expression of P-Selectin in HLECs to prevent CD4^+^ T cell differentiation (76). Further experiments are necessary to determine if the S1PR1/FOXC2 axis inhibits TLO formation by antagonizing LEC inflammation.

### Clinical relevance

IBD is a chronic inflammatory condition that affects the gastrointestinal tract and has substantial effects on all aspects of life (77). IBDs affects 1.4 million individuals in North America and 2.2 million individuals in Europe (78). The two main forms of IBD are ulcerative colitis and Crohn’s disease, both of which have an incidence of 3-20 per 100,000 in the United States (79). Crohn’s disease is treated with anti-inflammatory and immunosuppressive drugs, such as anti-TNF therapies, antibiotics, and surgery (80). These treatments manage symptoms, but never cure the disease, which causes episodic life-long illness. Thus, a better understanding of the etiopathology and mechanisms of Crohn’s disease is needed for the development of novel therapeutics (79). The presence of TLOs in the terminal ileum is one of the defining characteristics of Crohn’s disease (12, 81–84). Importantly, TLOs are thought to perpetuate inflammation by increasing immune cell retention and exacerbate tissue damage (81). Our discovery that *S1pr1^iΔLEC^* mice spontaneously develop TLOs in the terminal ileum without obvious tissue damage and inflammation appears to challenge this paradigm.

*Tnf^+/ΔARE^* mice weigh significantly less than their control littermates, have dysplastic intestinal epithelium and reduced survival (11, 26). In contrast, despite the presence of TLOs, *S1pr1^iΔLEC^* mice do not display changes in body weight, epithelial morphology or overall survival. It is possible that the TLOs in *Tnf^+/ΔARE^* mice are phenotypically distinct and more inflammatory when compared to those that are observed in *S1pr1^iΔLEC^*mice. The systemic effects of TNFα overexpression in *Tnf^+/ΔARE^* mice, such as poorly developed adipose tissue, cachexia and their potential effects on the intestinal epithelium may also contribute to the phenotype (85). Thus, the *S1pr1^iΔLEC^* mice could provide a complimentary model to understand the significance of TLOs in Crohn’s disease. We speculate that the TLOs in *S1pr1^iΔLEC^* mice are a sign of subclinical inflammation that does not cause tissue damage. Inflammatory triggers such as bacterial infections, food allergens or risk alleles might trigger an aggravated response by the immune cells in the TLOs resulting in IBD-like phenotype.

Finally, TLOs are important in the pathophysiology of several autoimmune diseases and cancer. Hence, it will be important to determine if *S1pr1^iΔLEC^*mice can develop TLOs in other organs such as the lungs if appropriate antigens, such as during influenza or COVID-19 infection or chronic smoking, become available. Whether tumor antigens can trigger TLO formation in *S1pr1^iΔLEC^*mice should also be tested. The effect of mesenteric tissue TLOs on metabolic disorders such as obesity should also be investigated. Nearly a dozen FDA-approved drugs are used for inhibiting S1PR1 and treating a variety of autoimmune diseases including multiple sclerosis and IBD (37). Our finding that the deletion of S1PR1 in LECs can result in the formation of TLOs raises both hope and concern regarding these drugs. On the one hand these drugs could be repurposed to trigger the formation of TLOs and potentiate immune checkpoint therapies for cancer. On the other hand, the formation of TLOs might compromise and even reverse the effectiveness of these drugs in treating autoimmune diseases. These possibilities must be investigated carefully.

## METHODS

### Mice

*Lyve1-Cre* (22)*, S1pr1^flox^* (41), S1PR1-GS (40) mice were described previously and were purchased from Jackson laboratory (catalog numbers 012601, 019141 and 026275 respectively)*. Sphk1^flox^*and *Sphk2^+/-^* mice were reported previously (22). Tg(Prox1-CreERT2) were a gift from Dr. Taija Makinen (Uppsala University) (30). *Vegfr3^+/EGFP^* mice were a gift from Dr. Hirotake Ichise (University of Ryukyus) (86). Mice were maintained in C57BL6 or C57BL6/NMRI mixed backgrounds and were fed standard chow diet. Tamoxifen was used for deleting S1PR1 in a time-specific manner. For early postnatal deletion, P1 S1pr1^iΔLEC^ and control littermate pups were fed 1 μl of 20 mg/ml tamoxifen, P3 pups 3 μl and so on until P7. For adult deletion, 8-wk-old mice were treated with tamoxifen by oral gavage (100 μg per gram of body weight) for three consecutive days. Tamoxifen stock was prepared by dissolving 200 mg tamoxifen (T5648, Sigma Aldrich Marketing Inc, St. Louis, MO, USA) in 10 ml of peanut oil and filter sterilized.

### Antibodies

The details about the antibodies that were used for IHC, western blotting and flow cytometry are provided as tables in the Supplementary Materials file.

### Cells

HLECs were a gift of Dr. Donwong Choi and Young-Kwon Hong (University of Southern California) (36, 87, 88). HLECs were grown on culture dishes or glass slides coated with 0.2% gelatin and were maintained in EGM-2 EC Growth Medium-2 Bullet Kit (Lonza). All experiments were conducted using cells until passage (P) 8. HLECs were treated as potential biohazards and were handled according to institutional biosafety regulations.

### Cell treatments and Analysis

#### siRNA transfection

HLECs were seeded onto modified culture dish as described previously (36, 89). Briefly, a 6-cm cell culture dish was adhered in the center of a 10-cm dish using non-toxic medical grade silicone glue (A-100, medical silicone adhesive, Factor II Inc, Lakeside, AZ, USA), UV irradiated for 1 hour and dried overnight. HLECs were seeded in the donut-shaped track. After 24 hours of culture (around 30% confluency), cells were transfected with siCTR (Cat# 51-01-14-03, Integrated DNA Technologies) or siS1PR1 (Cat# SI00376201, Qiagen) using Lipofectamine RNAiMax Transfection Reagent (Cat# 13778150, Invitrogen) according to manufacturer’s instruction.

#### Oscillatory Shear Stress

HLECs were seeded in culture dishes, transfected with siRNA as described above and grown to 80-100% confluency before exposing them to OSS for either 10 minutes or 24 hours. OSS was applied to the cells at ∼6 dynes/cm^2^ using the approach described previously (36). Briefly, the culture dishes were placed on top of bidirectional shaker (MS-NOR-30, Major Science) and horizontally rotated clockwise for ∼1 sec followed by anti-clockwise rotation for ∼1 sec at 100 rpm for 24 hours.

#### Cell shape and cell junction analysis

The ratio of cell length to width was calculated using NIS-Elements software (Nikon). For junction analysis, thick junction with two visible borders was identified as overlapping junctions (62). The ratio of the length of overlapping junction and that of total junction was quantified using NIS-Elements software (Nikon).

### Live vessel imaging and whole-mount immunofluorescence of isolated mesenteric lymphatics

Immediately after undergoing valve functional tests, 3-mo *S1pr1^f/f^;*Prox1-tdTomato (TM P1-7) and *S1pr1^iΔLEC^*; Prox1-tdTomato (TM P1-7) vessels were utilized for live confocal imaging of the tested lymphatic secondary valve. The isolated and cleaned mesenteric lymphatic vessels were cannulated in “Calcium Free” Krebs buffer to prevent movement artifacts during imaging. The vessels were imaged with an Andor Dragonfly 200 spinning disc confocal microscope on a Leica DMi8 inverted microscope equipped with a Zyla 4.2 Megapixel sCMOS camera and controlled by the Andor Fusion® Software. Image stacks of lymphatic valves, valve sites, or single valve leaflets were collected at 40X magnification (HCX PL APO 40x/1.10 W CORR) at 0.24-micron z-plane intervals throughout the entire vessel. 3D reconstructions were made using IMARIS software.

Following the valve functional test and live confocal imaging 3-mo *S1pr1^f/f^;*Prox1-tdTomato (TM P1-7) and *S1pr1^iΔLEC^*; Prox1-tdTomato (TM P1-7) vessels were used for immunostaining. While the vessels were still cannulated and pressurized, they were fixed using 1% chilled PFA for 20 minutes to maintain the vessel’s natural open state. Vessels were then fixed overnight in a 24 well dish on a rocker at 4C in 1%PFA and washed with PBS for 2-4 hours the following morning. The fixed mesenteric lymphatic vessels were permeabilized with PBS supplemented with 0.1 % TritonX100 for 30 minutes and then blocked in Blockaid buffer ® for 3 hours. The vessels were stained with primary and secondary antibodies overnight. Vessels were washed a final time in PBS and incubated with NucBlue (ThermoFisher; R37605 Hoechst 33342) in PBS for 5 minutes to stain the nuclei. Vessels were then canulated and imaged on the Andor Dragonfly using 40X magnification, 0.24-micron step size, and a Zyla 4.2 Megapixel sCMOS camera. Image stacks were then opened in IMARIS to create a 3D reconstruction for valve visualization. The “Crop 3D” tool was used to further restrict viewing to a single lymphatic valve leaflet. Display min max values and gamma values were optimized to assist in visualizing the fluorescent signal throughout the full image stack of the vessel.

### Multiplex enzyme-linked immunosorbent assay for cytokine

Serum cytokines were quantified using MILLIPLEX Mouse Cytokine/Chemokine Magnetic Bead Panel (EMD Millipore, Billerica, MA) and sample acquisition was performed on Bio-Plex 200 (BioRad, Hercules, CA) as per manufacturer’s instruction (90).

### Immunohistochemistry

IHC on cryosections, vibratome sections, mesentery and skin were performed according to our previously published protocols (29, 35, 69). Immunocytochemistry was performed as we described previously (35, 39, 91, 92).

For harvesting the adult mesentery, mice were euthanized by asphyxiation followed by perfusion with PBS and 4% PFA. Subsequently, the entire gut (stomach to rectum) was dissected and further fixed overnight in 2% PFA in the cold room. After washing profusely with PBS on ice, the mesenteries were dissected out from the gut and used for whole mount IHC using the iDISCO protocol that we described previously (92) with minor modifications. Specifically, the tissues were post-fixed in 4% PFA overnight in the cold room, washed with PBS and cleared for at least 3 days in the cold room with 1.62 M Histodenz medium and 0.1% Tween 20 (both from Sigma-Aldrich, St. Louis, MO, USA) (93). Cleared tissues were mounted on slides with 1.62 M Histodenz medium and images using LSM 710 laser-scanning microscope (Zeiss, Oberkochen, Germany) or C2+ confocal (Nikon, Tokyo, Japan) microscopes.

### Lymphangiography

Mice were anesthetized with ketamine/xylazine (25/2.5 mg/kg, i.p.) and placed face up on a heated tissue dissection/isolation pad. The abdomen was opened and a 2-3” section containing the cecum and terminal ileum was pulled out and pinned, with blood supply intact, onto a semi-circular base of Sylgard on the top of a transparent, water-jacketed base. The base was heated to 37°C. The preparation was continuously superfused with Krebs solution preheated to 37°C through a heating coil. The intestinal preparation was transferred to the stage of a Zeiss Axio-zoom V16 microscope for fluorescence imaging using Zeiss Zen software. The lymphatic networks in various regions of the terminal ileum were perfused with Krebs containing 1-2% FITC-dextran (Sigma) using sharpened micropipettes (∼40 um tip diameter). The micropipette was positioned using a Narishige micromanipulator such that it just penetrated the intestinal wall, usually adjacent to a pin in a Peyer’s patch. The pipette was pressurized from a 10 ml glass syringe as needed to perfuse the intestinal wall lymphatic capillary network. Once that network was perfused (usually for a length of intestinal wall of 1-2 mm), pressure was maintained for 1-30 min as the dye drained into the lymphatic collectors of the mesentery. Dye perfusion was monitored by continuously recording the FITC/GFP channel of the microscope at 5 fps. Once the main collector in that region was fully perfused, or after 20-30 min had elapsed without good collector perfusion (in the case of regions with TLOs), the pipette was withdrawn and the adjacent segment (1-2 mm away) was studied after a new micropuncture site was located. The typical procedure was to start imaging/perfusion at the last vascular arcade in the terminal ileum and then move proximally to perfuse the second and third arcades from the cecum.

#### Krebs solution contained

146.9 mM NaCl, 4.7 mM KCl, 2 mM CaCl_2_·2H_2_O, 1.2 mM MgSO_4_, 1.2 mM NaH_2_PO_4_·H_2_O, 3 mM NaHCO_3_, 1.5 mM Na-HEPES, and 5 mM D-glucose (pH = 7.4), supplemented with 0.5% bovine serum albumin for isolated vessel studies.

### Protein isolation and analysis

Protein was extracted from cells by using RIPA lysis buffer. Western blots were performed according to standard protocols. The intensities of bands were measured using ImageJ software.

### SEM

SEM was performed according to our previous protocol (29, 94). Briefly, vibratome sections were used for immunohistochemistry and confocal microscopy analyses. Subsequently, the tissues were fixed in 2% glutaraldehyde in 0.1 M cacodylate buffer for 2 hours. After washing profusely in PBS, the sections were post fixed in 1% osmium tetroxide in 0.1 M cacodylate buffer for 2 hours and subsequently dehydrated sequentially with increasing concentration of ethanol. The sections were further dehydrated in hexamethldisilazane (HMDS) and allowed to air-dry overnight. Dry sections were sputter-coated with Au/Pd particles (Med-010 Sputter Coater by Balzers-Union, USA) and observed under Quanta SEM (FEI, Hillsboro, OR, USA) at an accelerating voltage of 20KV.

### Valve function tests

The methods for assessing back-leak in isolated lymphatic collectors containing a single valve have been described in detail previously (72, 95) and documented in several recent studies of transgenic mice (42, 62, 96, 97). Mice were anesthetized with ketamine/xylazine (25/2.5 mg/kg, i.p.) and placed in the prone position on a heated tissue dissection/isolation pad. Mesenteric collectors were isolated by opening the abdomen, removing the entire small intestine and pinning it in a Sylgard-coated dish. Individual collectors were identified, excised and pinned to the chamber using 40 μm wire. After removing the majority of the associated fat and connective tissue, vessels containing a single valve were then transferred to a 3-ml myograph chamber containing Krebs-albumin solution and cannulated at each end with a glass micropipette (40-50 μm OD tip), pressurized slightly and further cleaned. The chamber with attached micropipettes, pipette holders and micromanipulators was transferred to the stage of an inverted microscope. Polyethylene tubing connected the back of each micropipette to low pressure transducers and a computerized pressure controller, allowing independent control of inflow (Pin) and outflow (Pout) pressures.

Valve function tests measured the pressure back-leak through a closed valve. Starting with Pin and Pout = 0.5 cmH_2_O, and the valve open, Pout was raised to 10 cmH_2_O, ramp-wise, over a 30-sec period while Pin was held at 0.5 cmH_2_O. Normal valves closed as Pout exceeded ∼1 cmH_2_O and remained closed for the duration of the Pout ramp. In some cases when valves appeared stiffer than normal, gentle tapping of the Pout line was used to encourage closure. Pressure back-leak through the closed valve was measured with a servo-null micropipette inserted through the vessel wall on the inflow side of the vessel, which could resolve changes as small as ∼0.05 cmH_2_O. The value of Psn at the end of the ramp (Pout = 10 cmH_2_O) was used as the standard index of back-leak.

### Statistics

For biochemical studies the number n refers to the number of times the experiment was performed. For histochemical analysis n refers to the total number of animals included per group. Statistically significant differences were determined using unpaired *t* test, Mann-Whitney test or 2-way ANOVA with Tukey’s post-hoc tests. Prism software was used for statistical analyses. Data are reported as mean ± SD with significance set at p < 0.05. n and p values for each experiment is provided in the figure legends. Western blots are performed at least three independent times. The most representative Western blots are presented.

### Study Approval

All mice were housed and handled according to the institutional IACUC protocols.

## AUTHOR CONTRIBUTIONS

XG, MJD and RSS designed and performed experiments, analyzed the results and wrote the manuscript; LC, ZA, GPF, YH, IDG, MND, XS, RSK, and SZ performed experiments and analyzed the results; HC, FL, LX, GJR and EC provided critical input and reagents; all authors provided input in editing the manuscript.

## ACKNOWLEDGEMENTS

We thank Dr. Taija Makinen (Uppsala University) and Dr. Hirotake Ichise (University of Ryukyus) for Tg(Prox1-CreERT2) and *Vegfr3^+/GFP^* mouse lines respectively. We thank Dr. Donwong Choi and Young-Kwon Hong (University of Southern California) for HLECs, Ms. Lisa Whitworth (Microscopy Laboratory, Oklahoma State University, Stillwater) for SEM, and Dr. Lorin E. Olson (OMRF) for insightful comments. This work is supported by NIH/NHLBI (R01HL131652 and R01HL163095 to RSS; R01HL133216 to RSS and HC; R01HL122578 to MJD; R01HL175083 to SDZ), Oklahoma Center for Adult Stem Cell Research, a program of TSET (4340) to RSS, NIH/NIGMS COBRE (P20 GM139763 to LX), and the French National Research Agency (ANR-19-CE14-0028-01) to EC.

## Non-standard abbreviations

LECs: lymphatic endothelial cells

LVs: lymphatic valves

LVVs: lymphovenous valves

HLEC: primary human LECs

OSS: Oscillatory shear stress

TLO: tertiary lymphoid organs.

