## Supplemental Figures and Legends for "S1PR1 Regulates Lymphatic Valve Development And Prevents Ileitis-Independent Tertiary Lymphoid Organ Formation"

#### Supplementary Materials

##### Antibodies

Antibodies that were used for immunohistochemistry.

| Antibody | Manufacturer | Catalog Number |
|---|---|---|
| GFP | abcam | ab13970 |
| VEGFR3 | R&D SYSTEMS | AF743 |
| CD31 | BD Pharmingen | 553370 |
| PROX1 | AngioBio Co. | 11-002 |
| CLAUDIN5 | Thermo Fisher Scientific | 34-1600 |
| LYVE1 | R&D SYSTEMS | AF2125 |
| B220 | Biolegend | 13204 |
| S100A9 | R&D SYSTEMS | AF2065 |
| RFP | Rockland | 200-101-379 |
| CD11c | Biolegend | 117301 |
| GL7 | Invitrogen | 48-5902-82 |
| F4/80 | Abcam | ab6640 |
| CCL21 | R&D SYSTEMS | AF366 |
| CD3e | Invitrogen | 16-0031-85 |
| CD11b | Invitrogen | 14-0112-82 |
| ICAM1 | R&D SYSTEMS | AF796 |
| MADCAM1 | Millipore Sigma | MABF2072 |
| F-ACTIN | Invitrogen | A12379 |
| VE-CADHERIN | BD Pharmingen | 550548 |
| S1PR1 | R&D SYSTEMS | MAB7089 |
| FOXC2 | R&D SYSTEMS | AF6989 |
| GATA2 | R&D SYSTEMS | AF2046 |
| CX37 | Invitrogen | 40-4200 |
| ITGa9 | R&D SYSTEMS | AF3827 |
| Prox1 | R&D SYSTEMS | AF2727 |
| Cy3-conjugated Donkey Anti-Goat IgG (H+L) | Jackson ImmnunoReserach Lab | 705-165-147 |
| Cy3-conjugated Donkey Anti-Rabbit IgG (H+L) | Jackson ImmnunoReserach Lab | 711-165-152 |
| Cy5-conjugated Donkey Anti-Rat IgG (H+L) | Jackson ImmnunoReserach Lab | 712-175-150 |

| | | |
|---|---|---|
| Cy5-conjugated Donkey Anti-Goat IgG (H+L) | Jackson ImmnunoReserach Lab | 705-175-147 |
| Cy5-conjugated Donkey Anti-Sheep IgG (H+L) | Jackson ImmnunoReserach Lab | 713-175-147 |
| Alexa Fluor 488-conjugated Donkey Anti-Rabbit IgG (H+L) | Jackson ImmnunoReserach Lab | 711-547-003 |
| Alexa Fluor 488-conjugated Donkey Anti-Goat IgG (H+L) | Jackson ImmnunoReserach Lab | 705-545-147 |
| Alexa Fluor 488-conjugated Donkey Anti-Chicken IgG (H+L) | Jackson ImmnunoReserach Lab | 703-545-155 |
| Alexa Fluor 488-conjugated Donkey Anti-Rat IgG (H+L) | Jackson ImmnunoReserach Lab | 712-545-153 |
| Alexa Fluor 488-conjugated Goat Anti-Hamster IgG (H+L) | Invitrogen | A-21110 |

Antibodies that were used for whole-mount IHC of isolated mesenteric lymphatic vessels.

| Antibody | Manufacturer | Catalog Number |
|---|---|---|
| PROX1 | Abcam | ab101851 |
| CD31 | BD Pharmingen | 550274 |
| ITGa9 | R&D SYSTEMS | AF3827 |
| Alexa Fluor 488-conjugated donkey anti-rabbit | ThermoFisher | A-21206 |
| Alexa Fluor 555-conjugated donkey anti-rat | ThermoFisher | A78945 |
| Alexa Fluor 647-conjugated donkey ant-goat | ThermoFisher | A-21447 |

Antibodies that were used for western blotting.

| Antibody | Manufacturer | Catalog Number |
|---|---|---|
| FOXC2 | R&D SYSTEMS | AF5044 |
| FOXO1 | Cell Signaling Technology | CST2880 |
| pFOXO1 | Cell Signaling Technology | CST2486 |
| b-ACTIN | SIGMA-ALDRICH | A5441 |
| pAKT | Cell Signaling Technology | 4060 |
| AKT | Cell Signaling Technology | 4691 |
| pERK | Cell Signaling Technology | 4376 |
| ERK | Cell Signaling Technology | 4695 |
| GAPDH | Cell Signaling Technology | 5174 |

| | | |
|---|---|---|
| HRP-conjugated Goat anti-Rabbit IgG (H+L) | ImmunoReagents, Inc | GtxRb-003-EHRPX |
| HRP-conjugated Donkey anti-Sheep IgG | R&D SYSTEMS | HAF016 |

**Supplementary Figures and Figure Legends**

**Supplementary Figure 1**

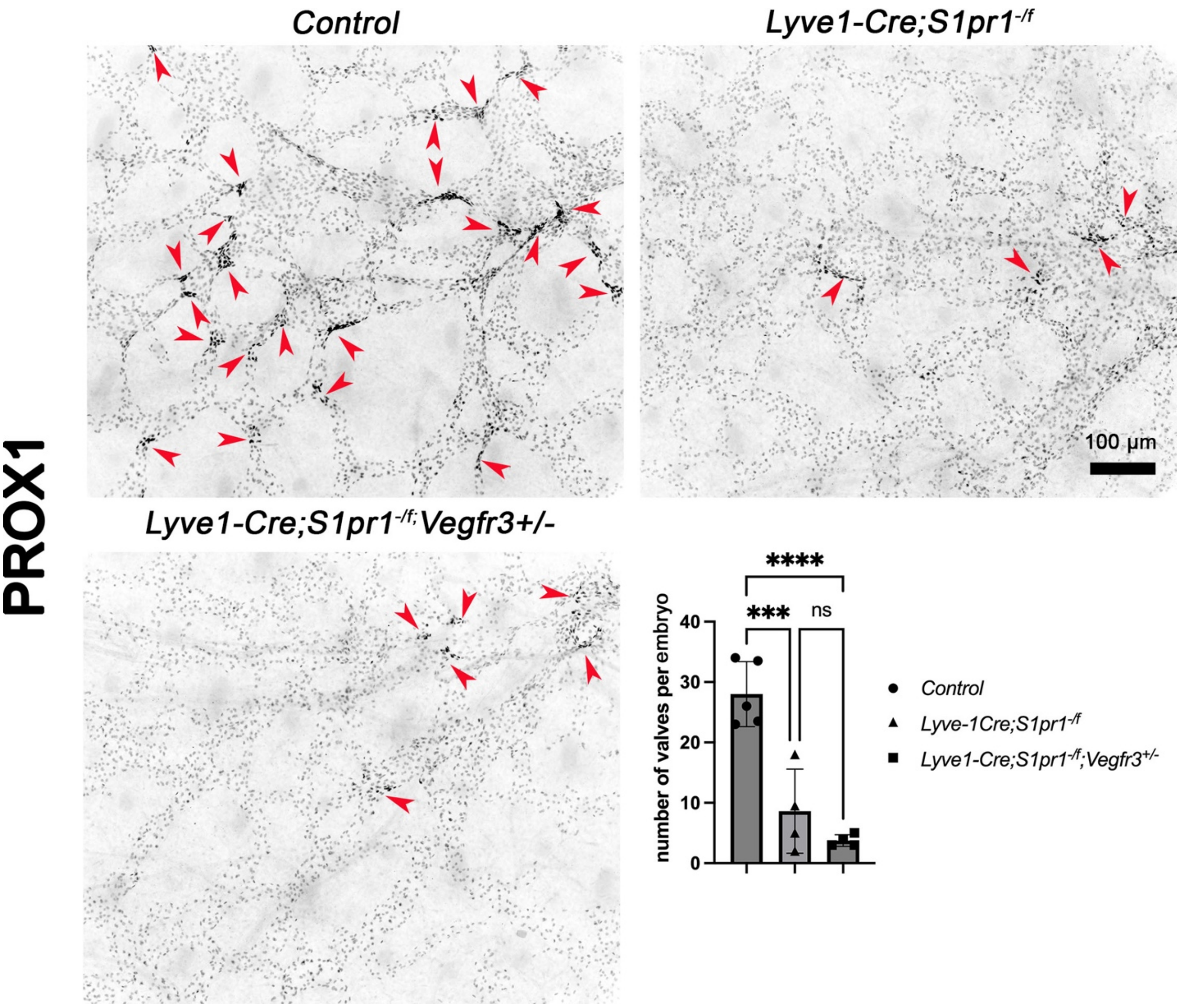


**Supplementary Figure 1: S1PR1 regulates dermal LV development in a VEGFR3-independent manner.**

The dorsal skin of E16.5 embryos were dissected and analyzed by whole mount immunohistochemistry with anti-PROX1 antibody. The developing LVs were identified by the presence of PROX1hi clusters (arrowheads).

Statistics: n=5 for controls embryos; n=4 for each mutant genotype respectively. Each dot represents one animal on the graph. The graphs were plotted as Mean ± SD. One way ANOVA was performed for the statistical analysis. *** $p<0.01$; **** $p<0.001$.

#### Supplementary Figure 2

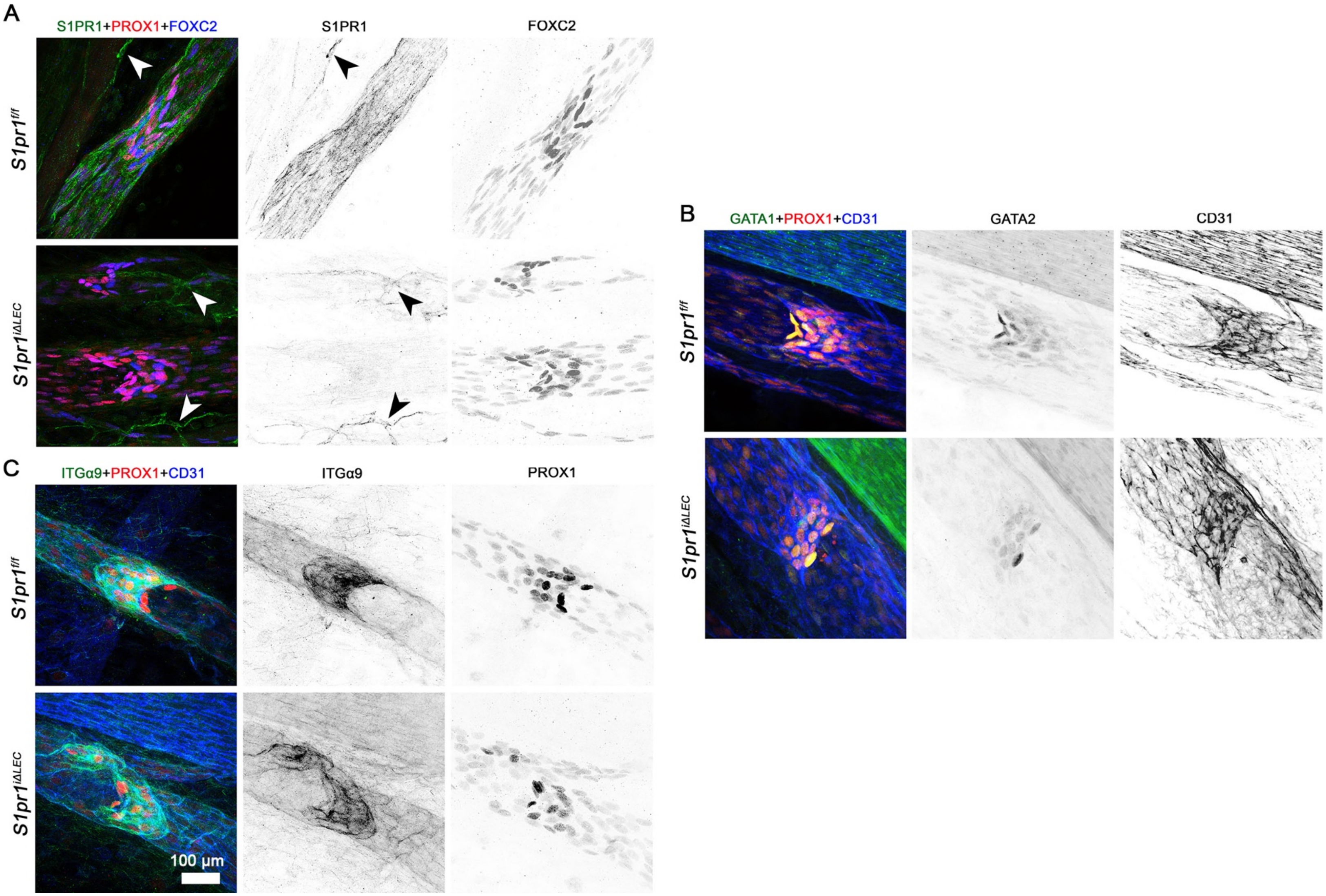

**Supplementary Figure 2: LVs remaining after efficient deletion of S1PR1 express valve markers normally.**

The mesenteric vasculature of P10 *S1pr1*$^{f/f}$ (TM @ P1-7) and *S1pr1*$^{i\Delta LEC}$ (TM @ P1-7) littermates were analyzed using the indicated antibodies. (A) IHC confirmed that S1PR1 was specifically and efficiently deleted from the lymphatic vessels and LVs of mutants. Arrowheads point to blood capillaries in which S1PR1 expression was comparable between the two genotypes. (A-C) The expression of valve markers PROX1 (A-C), FOXC2 (A), GATA2 (B), and Integrin-$\alpha$9 (C) appeared normal in the remaining LVs of mutants.

Statistics: n=3 per genotype.

**Supplementary Figure 3**

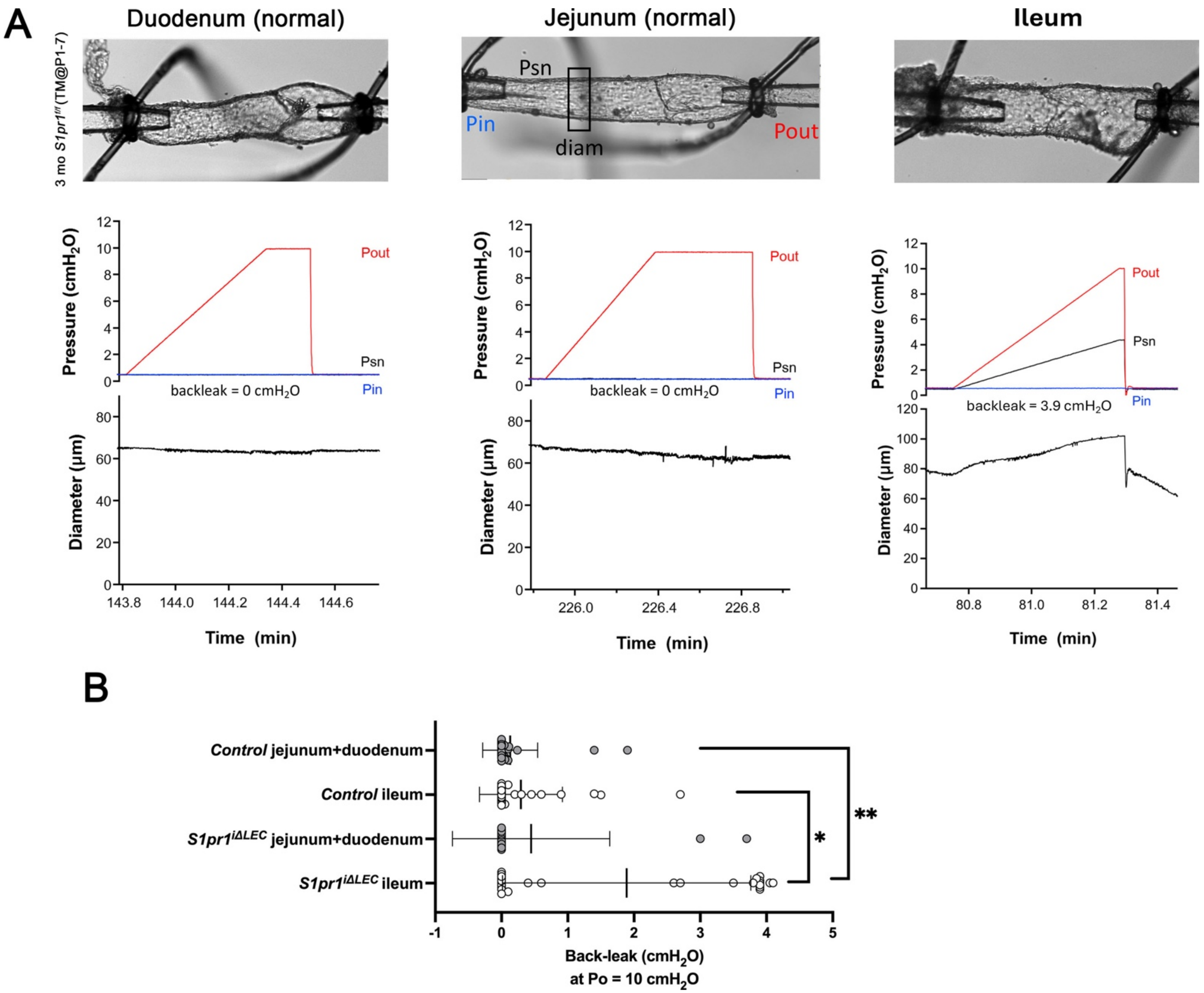

**Supplementary Figure 3: Most lymphatic valves located in the proximal mesenteric lymphatic vessels of *S1pr1$^{i\Delta LEC}$* mice were normal.**

(A) Mesenteric lymphatic vessels from the sections corresponding to the duodenum, jejunum and ileum were dissected from 3-month-old *S1pr1$^{i\Delta LEC}$* (TM @ P1-7) mice, cleaned and cannulated (top row). Subsequently, LV function test was performed using two approaches. While gradually increasing the outflow pressure (Pout) the pressure (Psn) and vessel diameter were measured behind the LVs. Duodenal and jejunal LVs did not have any back leak (second row). Consequently, the lymphatic vessels did not dilate (bottom row). In contrast, LVs from the ileum had back leak as indicated by increasing Psn and diameter with increasing Pout. (B) Quantification of back leak of LVs harvested from the duodenum and jejunum versus ileum of control and *S1pr1$^{i\Delta LEC}$* mice.

Statistics: B) n=31 control (duodenum and jejunum), n=28 control (ileum), n=15 *S1pr1$^{i\Delta LEC}$* (duodenum and jejunum) and n=26 *S1pr1$^{i\Delta LEC}$* (ileum) LVs. LVs from 3-mo *S1pr1$^{i\Delta LEC}$* (TM @ P1-P7) and 10-mo *S1pr1$^{i\Delta LEC}$* (TM @ 8w) were combined as *S1pr1$^{i\Delta LEC}$* for this analysis.

**Supplementary Figure 4**

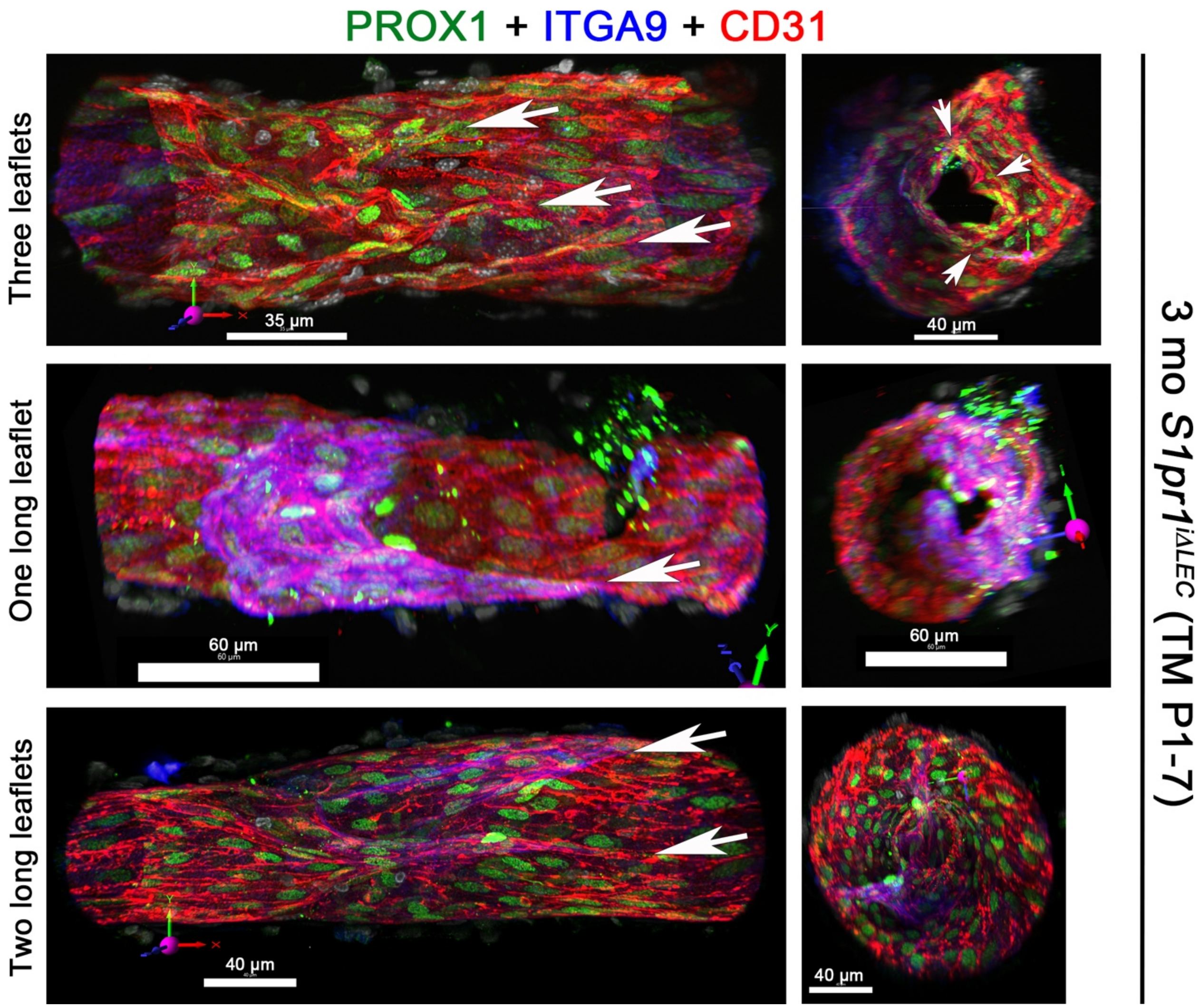


**Supplementary Figure 4: Heterogeneous LV defect in *S1pr1$^{i\Delta LEC}$* mice.**

Lymphatic vessels with leaky LVs from 3 mo *S1pr1$^{i\Delta LEC}$* (TM@P1-7) mice were fixed and whole-mount IHC was performed for the indicated markers. Confocal imaging and 3D reconstruction were performed to identify the structural defects in the LVs. One LV appeared to have 3 leaflets (arrows). One or both leaflets were abnormally elongated at their insertion points in two other LVs resulting in abnormal *en face* LV structure (right).

Statistics: n=5 leaky LVs were analyzed.

**Supplementary Figure 5**

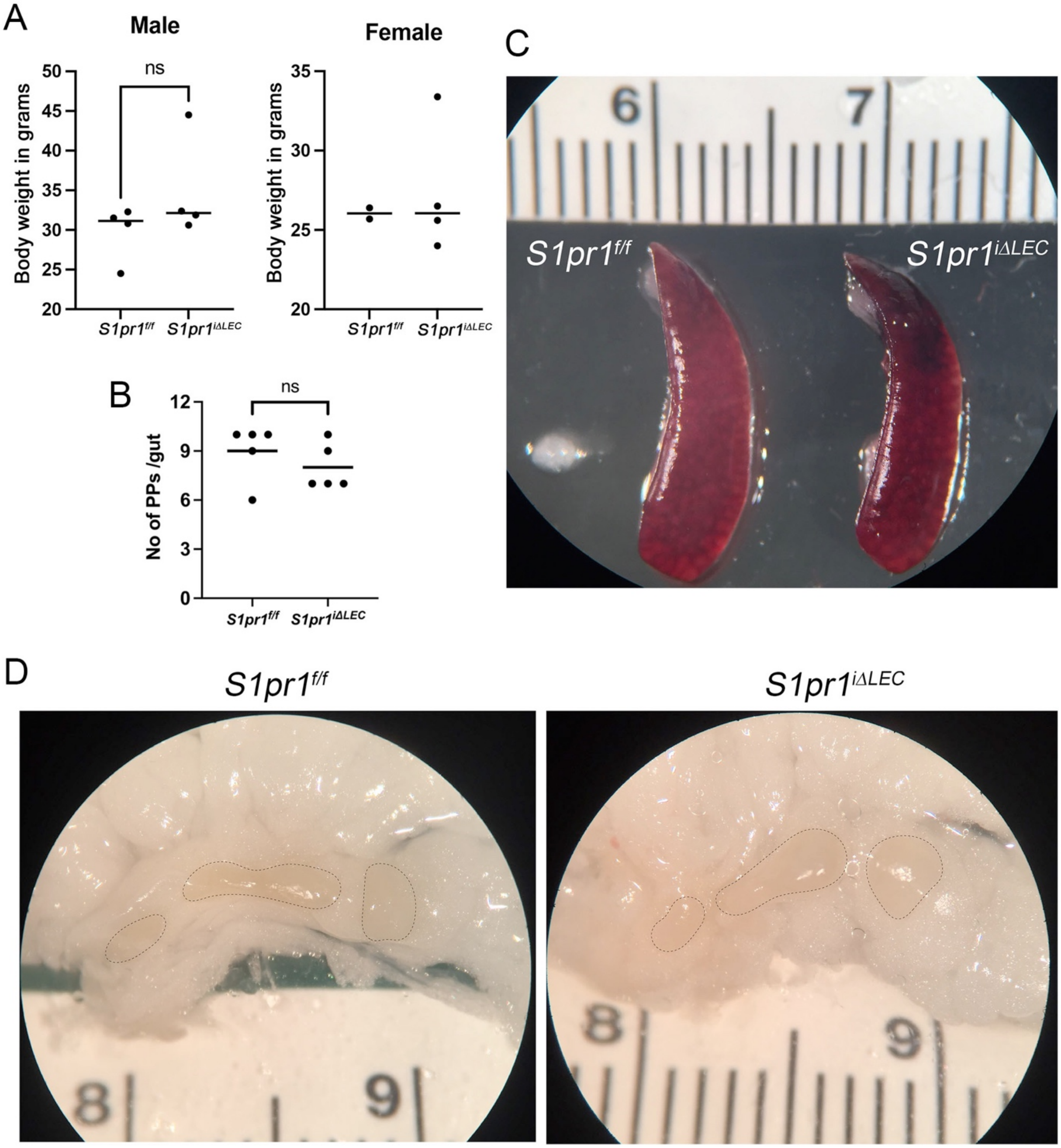

**Supplementary Figure 5: *S1pr1*$^{i\Delta LEC}$ mice did not have obvious characteristics of inflammatory disease.**

(A) 10-month-old *S1pr1*$^{f/f}$ (TM @ P1-7) and *S1pr1*$^{i\Delta LEC}$ (TM @ P1-7) littermates had comparable body weights irrespective of sex.

(B) The number of Payer's patches in the guts (duodenum to cecum) of 10-month-old *S1pr1*$^{f/f}$ (TM @ P1-7) and *S1pr1*$^{i\Delta LEC}$ (TM @ P1-7) littermates was counted and found to be comparable.

(C, D) Spleen (C) and mesenteric lymph nodes (D) of 10-month-old *S1pr1*$^{f/f}$ (TM @ P1-7) and *S1pr1*$^{i\Delta LEC}$ (TM @ P1-7) littermates were comparable in size and shape.

Statistics: (A) Each dot in the graph represents an individual animal. n=4 male *S1pr1*$^{f/f}$, n=4 male *S1pr1*$^{i\Delta LEC}$, n=2 female *S1pr1*$^{f/f}$ and, n=4 female *S1pr1*$^{i\Delta LEC}$. Statistical significance was calculated for males using Mann-Whitney test; (B) n=4 males and 1 female per genotype. Statistical significance was calculated for males using Mann-Whitney test; (C, D) Images are representative of n=3 animals/genotype/sex.

### Supplementary Figure 6

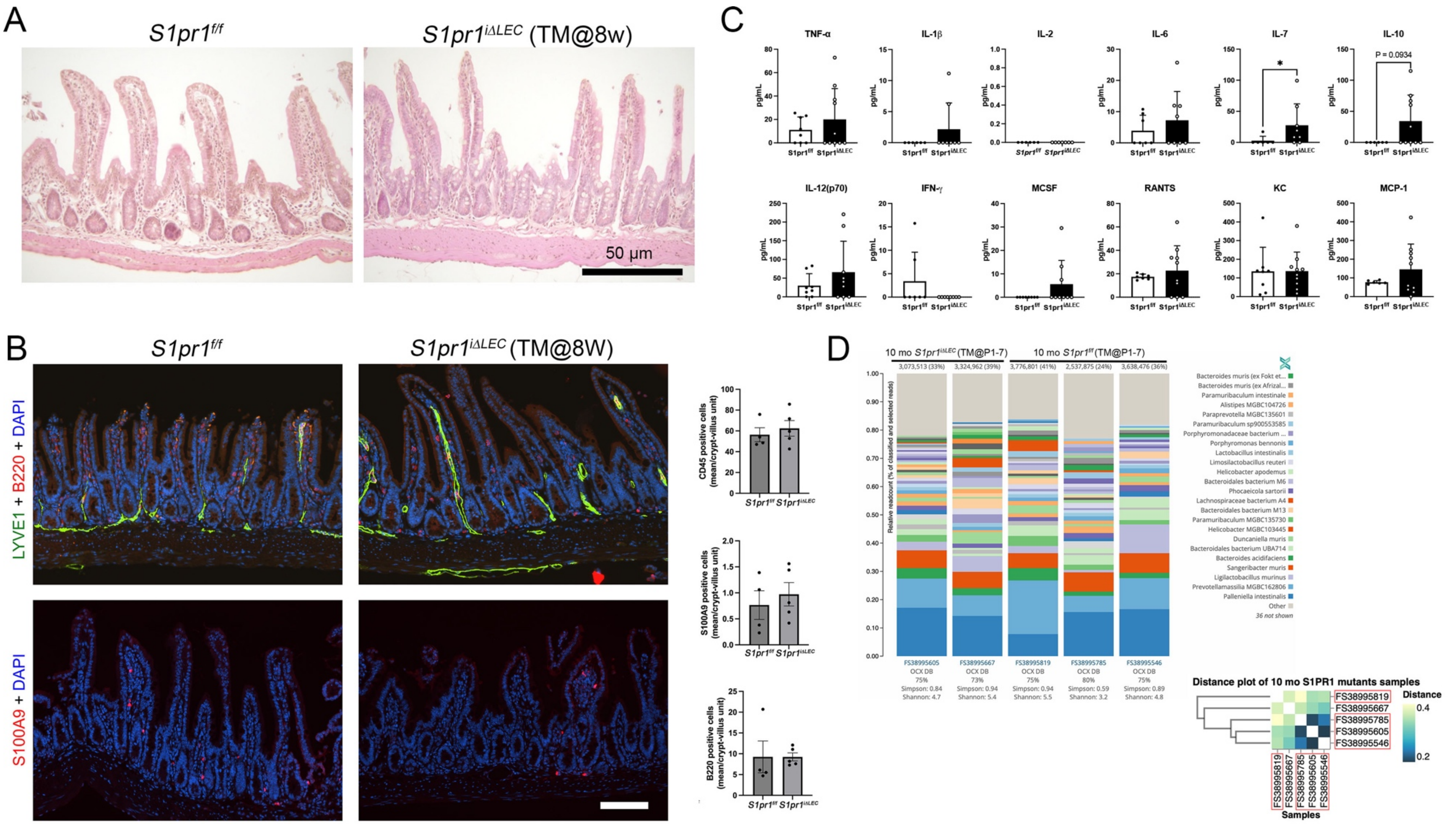

**Supplementary Figure 6: *S1pr1$^{i\Delta LEC}$* mice do not develop epithelial dysplasia, ileal inflammation or microbial dysbiosis.**

(A, B) The ileum of *S1pr1$^{f/f}$* and *S1pr1$^{i\Delta LEC}$* mice (TM@8w) were sectioned and analyzed by H&E (A) or IHC for the indicated markers (B). (A) The intestinal epithelium of *S1pr1$^{i\Delta LEC}$* mice appeared to be indistinguishable from control samples with no obvious infiltration of immune cells. (B) The same number of CD45$^+$ hematopoietic cells, B220$^+$ B cells and S100A9$^+$ neutrophils were observed in the control and *S1pr1$^{i\Delta LEC}$* mice.
(C) The serum from *S1pr1$^{f/f}$* and *S1pr1$^{i\Delta LEC}$* mice was analyzed by multiplex ELISA for inflammatory cytokines. IL-7 was modestly increased in the *S1pr1$^{i\Delta LEC}$* mice. No significant differences were observed between the control and *S1pr1$^{i\Delta LEC}$* mice for the other cytokines.
(D) The fecal pellets of *S1pr1$^{f/f}$* and *S1pr1$^{i\Delta LEC}$* mice were analyzed by shotgun metagenomic analysis. The numbers on top of the graphs indicate the number of bacterial reads. The same information is provided within brackets as a percentage of total reads. The graphs indicate the relative abundance of the bacterial species. No obvious change was observed between the *S1pr1$^{f/f}$* and *S1pr1$^{i\Delta LEC}$* mice. The inset is a hierarchical clustering of the same samples with the co-housed mice within red boxes. Three of the four co-housed mice appear to be closer to each other irrespective of their genotype.

Statistics: (A, B) n=4 control and n=5 *S1pr1$^{i\Delta LEC}$* mice (TM@8w). 2-4 sections per mice were analyzed and the number of cells per crypt-villus unit was calculated and graphs were plotted as mean $\pm$ SD. Normality was tested using Shapiro-Wilk method, variance was determined by F test. Significance was determined using unpaired t test for CD45 and S100A9, and Mann-Whitney test for B220; (C) n=8 control and n=10 *S1pr1$^{i\Delta LEC}$* mice (TM@8w). Graphs were plotted as mean $\pm$ SD. Normality was tested using Shapiro-Wilk method, variance was determined by F test. Significance was determined using unpaired t test with Welch's correction for MCP-1 and RANTS, and Mann-Whitney test for the other cytokines. (D) n=2 male *S1pr1$^{i\Delta LEC}$* (TM@P1-7) and n=3 tamoxifen treated *S1pr1$^{f/f}$* control littermates. In the distance plot cohoused mice are within the red box.

**Supplementary Figure 7:**

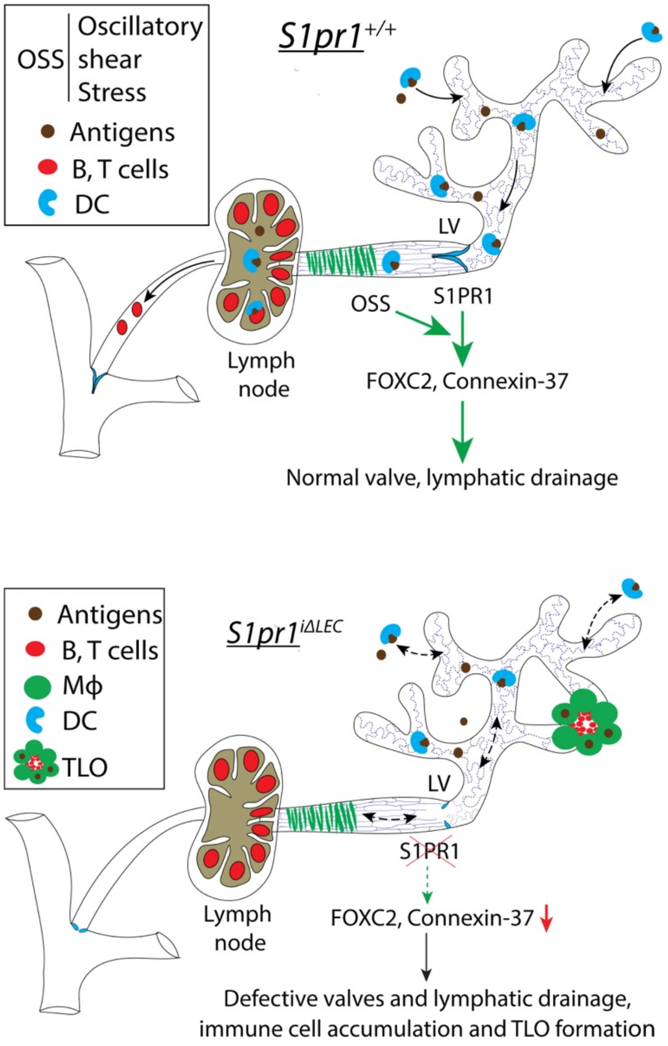


**Supplementary Figure 7: Working model**
LVs are necessary for proper immune cell drainage to the lymph nodes. S1PR1 regulates LV development by enhancing the expression of FOXC2 and Connexin-37 in response to OSS. LV defects caused by the loss of S1PR1 results in immune cell accumulation within lymphatic vessels and the mesenteric adipose tissue, chronic low-grade inflammation and TLO formation.

**Additional Supplementary Files**

1. Supplementary Movie 1: Live imaging of fluorescent dye flow along the ileal lymphatic vessels of a 1-year-old *S1pr1*$^{f/f}$ mouse.
2. Supplementary Movie 2: Live imaging of fluorescent dye flow along the ileal lymphatic vessels of a 1-year-old *S1pr1*$^{i\Delta LEC}$ mouse (sample 1).
3. Supplementary Movie 3: Live imaging of fluorescent dye flow along the ileal lymphatic vessels of a 1-year-old *S1pr1*$^{i\Delta LEC}$ mouse (sample 2).
